# Bayesian test of gene flow between sister lineages using genomic data

**DOI:** 10.1101/2025.08.31.673316

**Authors:** Ziheng Yang, Xiyun Jiao, Sirui Cheng, Tianqi Zhu

## Abstract

Inference of interspecific gene flow using genomic data is important to reliable reconstruction of species phylogenies and to our understanding of the speciation process. Gene flow is harder to detect if it involves sister lineages than nonsisters; for example, most heuristic methods based on data summaries are unable to infer gene flow between sisters. Likelihood-based methods can identify introgression between sisters but the test exhibits several nonstandard features, including boundary problems, indeterminate parameters, and multiple routes from the alternative to the null hypotheses. In the Bayesian test, those irregularities pose challenges to the use of the Savage-Dickey (S-D) density ratio to calculate the Bayes factor. Here we develop a theory for applying the S-D approach under nonstandard conditions. We show that the Bayesian test of introgression between sister lineages has low false-positive rates and high power. We discuss issues surrounding the estimation of the rate of gene flow between sister lineages, especially at very low or very high rates, and suggest that evidence for gene flow between sisters be assessed via a Bayesian test. We find that the species split time has a major impact on the information content in the data, with more information at deeper divergence. We use a genomic dataset from *Sceloporus* lizards to illustrate the test of gene flow between sister lineages.

## Introduction

### Genomic data can be used to test introgression between sister species

Detecting gene flow between species and estimating the rate of gene flow is important to reconstructing species phylogenies and testing theories of speciation (Sousa and Hey, 2013; Payseur and Rieseberg, 2016; Roux *et al*., 2016). A number of methods have been developed to infer gene flow using genomic data, including heuristic methods based on data summaries and likelihood methods based on parametric models of gene flow (Jiao *et al*., 2021; Hibbins and Hahn, 2022). The former class of methods includes the *D*-statistic (or ABBA-BABA test) (Green *et al*., 2010), HyDe (Blischak *et al*., 2018) and PhyNest (Kong *et al*., 2025), which use genome-wide site-pattern counts in a species quartet; SNaQ (Solis- Lemus and Ane, 2016; Solis-Lemus *et al*., 2017; see also Yu *et al*., 2012) which uses reconstructed gene tree topologies; and *∂*a*∂*i (Gutenkunst *et al*., 2009) and fast-simcoal2 (Excoffier *et al*., 2013), which use the joint site frequency spectrum at single nucleotide polymorphism (SNP) sites. The latter class of methods, including both maximum likelihood (ML) and Bayesian inference, has the major feature that they use the multispecies coalescent (MSC) model with gene flow to average over genealogical histories underlying sequences sampled at each locus (Jiao *et al*., 2021). In ML, this is achieved through numerical integration (Zhu and Yang, 2012; Dalquen *et al*., 2017), while in Bayesian inference, it is through Markov chain Monte Carlo (MCMC) algorithms. Gene flow has been modelled as either a discrete introgression/hybridization event which occurred at a particular time point in the past (the MSC-I model; e.g., PhyloNet, Wen and Nakhleh, 2018; bpp, Flouri *et al*., 2020) or as a continuous process with a constant migration rate per generation (the MSC-M model; IMa3, Hey *et al*., 2018; bpp, Flouri *et al*., 2023). Our main focus in this paper is inference under the discrete introgression model (MSC-I) in the Bayesian framework.

Introgression may be harder to detect if it involves sister lineages than nonsisters. For example, gene flow between nonsisters may cause asymmetries in the gene tree distribution and in site-pattern counts in genomic data for a species quartet, while gene flow between sisters does not. As a result, quartet summary methods such as *D*, HyDe and SNaQ, which rely on counts of estimated gene trees or site patterns for species quartets, can detect gene flow between nonsisters but not between sisters. Likelihood methods use information in the coalescent times or branch lengths on gene trees, and in the genealogical variation across the genome, and can thus infer gene flow between sister lineages (Jiao *et al*., 2021; Hibbins and Hahn, 2022). They also accommodate multiple samples from the same species (in particular from the recipient lineage), which may boost the information content in the data about gene flow considerably (Jiao *et al*., 2021; Thawornwattana *et al*., 2026). In simulations, Bayesian inference under both MSC-I and MSC-M models showed similar power in inferring gene flow between sisters to that between nonsisters (Huang *et al*., 2020; Thawornwattana *et al*., 2023, Thawornwattana *et al*., 2025).

We note that in this context “sister species” simply means that the lineages involved in introgressive hybridization are sister branches on the phylogeny, and does not even imply that the two species are closely related. For example, given the phylogeny (*A*, (*B, C*)) , the *A* → *B* introgression is between nonsisters, but is between sisters if species *C* is not included in the data (e.g., if *C* is extinct or unsampled). Thus the distinction between gene flow involving sisters and that involving nonsisters has no biological significance. Also in this paper we use the terms species, populations, and lineages interchangeably.

### The Bayes factor for testing nested hypotheses can be calculated via the Savage-Dickey (S-D) density ratio

The standard device for model comparison in the Bayesian framework is the Bayes factor, defined as the ratio of the marginal likelihood values between the two compared models (Jeffreys, 1935). The marginal likelihood under a model is an average (often a high-dimensional integral) over the parameters under the model. This typically requires expensive computation involving many MCMC runs (Fourment *et al*., 2020, see also Discussion). In the special case of comparing nested hypotheses, the Bayes factor may be calculated via the Savage-Dickey (S-D) density ratio (Dickey, 1971; Ji *et al*., 2023), which involves far less computation. In phylogenetics Suchard *et al*. (2003) used the S-D density ratio to conduct Bayesian tests of the molecular clock; however, in that study the phylogeny is allowed to differ between the clock and no-clock models which appears to invalidate the S-D approach.

Let *H*_0_ be the null hypothesis and *H*_1_ the alternative hypothesis, with *H*_0_ being a special case of *H*_1_. In other words, *H*_1_ reduces to *H*_0_ when certain parameters of interest in *H*_1_ take special values (called the *null value*): *ω* = *ω*_0_. Suppose ‘nuisance parameters’ *λ* exist in both models, so that the prior under *H*_0_ is *π*_0_ (*λ*) and that under *H*_1_ is *π* (*λ, ω*) = *π ω π* (*λ* | *ω*). On the condition that the priors on nuisance parameters match between the two models at the null value,

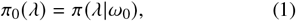

the Bayes factor *B*_10_ in support of *H*_1_ against *H*_0_ is given by

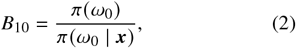

where ***x*** represent the data, and *π* (*ω*_0_) and *π* (*ω*_0_)| ***x*** are the prior and posterior densities of *ω*, respectively, under *H*_1_ evaluated at *ω* = *ω*_0_. Dickey (1971) provided a proof of eq. 2, which accounts for presence of nuisance parameters (*λ*) but does not accommodate the possible presence of unidentifiable parameters. This is dealt with in the extended proof of Ji *et al*. (2023).

### Irregularities of the test of introgression between sister lineages and objectives of this paper

Consider two species that diverged time *τ* ago with the phylogeny (*A, B*) (Fig. 1). In analysis of genomic sequence data, it is convenient to measure time by mutations, and *τ* is thus measured in the expected number of mutations per site. One time unit is then the expected time to accumulate one mutation per site. At this time scale, the coalescence rate for two sequences sampled from a diploid species with population size *N* is 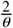 (i.e., 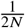 per generation), where *θ* = 4*N μ* is known as the population size parameter, with *μ* to be the mutation rate per site per generation. Also the ‘coalescent time unit’ or the expected waiting time for two sequences to coalesce is 2*N* generations or 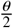 mutations per site. The MSC model with no gene flow is the null hypothesis (*H*_0_) with parameters *τ* and *θ* (Fig. 1A). In the alternative hypothesis (*H*_1_), introgression occurs from species *A* to *B* (i.e., from *X* to *Y*) at time *τ*_*X*_ ≡ *τ*_*Y*_ , with the introgression probability *φ* ≡ *φ*_*XY*_ defined as the proportion of immigrants in the recipient population *Y* from the donor population *X* (Fig. 1B). Here we let time run forward when we define parameters. Both *H*_0_ and *H*_1_ involve a population size parameter *θ*, which is dropped in our notation. The parameter vector is thus ***θ***_0_ = (*τ*) for *H*_0_ and ***θ*** = (*φ, τ*_*Y*_ , *τ*_*R*_) for *H*_1_ (Fig. 1A&B).

**Fig. 1:**
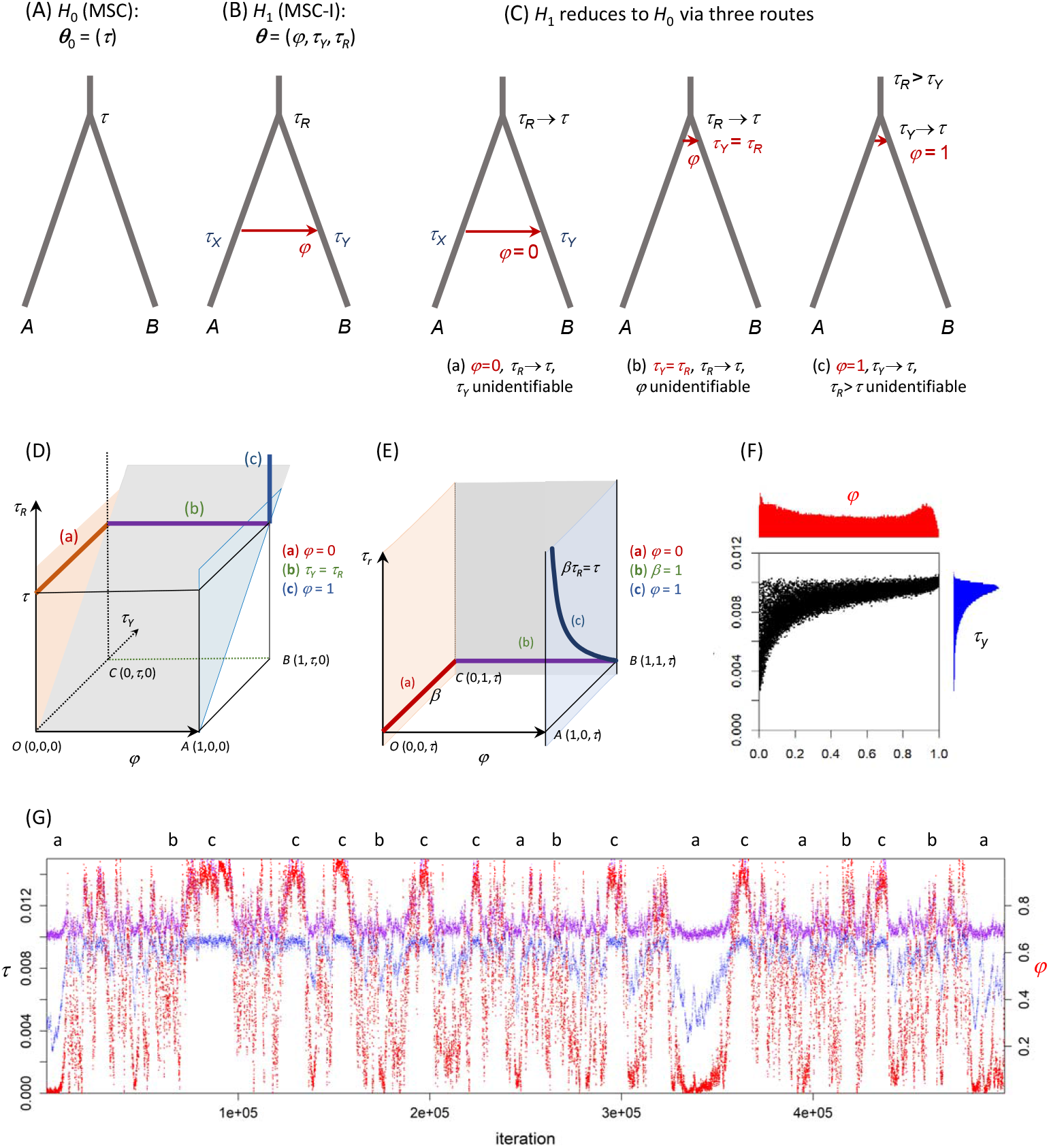
(**A, B**) The null (*H*_0_) and alternative (*H*_1_) hypotheses in the test of (unidirectional) introgression from *A* to *B*. The parameter vector is ***θ***_0_ = (*τ*) under *H*_0_ and ***θ*** = (*φ, τ*_*Y*_ , *τ*_*R*_) under *H*_1_, with the population size parameter (*θ*) suppressed. (**C**) *H*_1_ reduces to *H*_0_ via three routes (constraints): a, b, and c (table 1(i)). (**D**) Parameter space of *H*_1_, showing the null point as three lines a, b, c, which correspond to the three routes of panel **C**, and the null space (the parameter space for *H*_0_) as three planes. (**E**) Reparametrization of *H*_1_, with *β* = *τ*_*Y*_/*τ*_*R*_ used instead of *τ*_*Y*_ , so that ***θ***^′^ = (*φ, β, τ*_*R*_). (**F**) The 2-D scatter plot (for *τ*_*Y*_ and *φ*) and (**G**) trace plots for the three parameters in *H*_1_ from an MCMC sample generated in a bpp analysis of a dataset simulated under *H*_0_: purple for *τ*_*R*_, blue for *τ*_*Y*_ and red for *φ* (the 2nd *y*-axis). Labels on top of the trace plot (**G**) indicates that the chain is in the neighborhood of segments a, b, c of panel **D**. For example, close to segment a, *φ* ≈ 0, *τ*_*R*_ ≈ *τ*, while *τ*_*Y*_ is indeterminate. The dataset consists of 500 loci, with 10 sequences per species per locus, and with 500 sites in each sequence, simulated under *H*_0_ with *τ* = 0.01 and *θ* = 0.005. The data are analyzed under *H*_1_, using the priors *θ* ∼ *G* (2, 200) , *τ*_*R*_ ∼ *G* (2, 200) , and *φ* ∼ *U* (0, 1). The same data are analyzed using different priors on *φ* in Figure S3.

While *H*_0_ is clearly nested within *H*_1_, the test comparing the two models involves several non-standard features.

**Table 1:** Parameters of interest (*ω*), nuisance parameters (*λ*), and indeterminate parameters (*ξ*) and condition on priors in Bayesian test of introgression.

| Segment | Parameters of interest<br>($\omega = \omega_0$) | Indeterminate<br>($\xi$) | Nuisance<br>($\lambda$) | Condition<br>on $\lambda$ (eq. 5) |
| --- | --- | --- | --- | --- |
| **(i) Unidirectional introgression (UDI, Fig. 1C)** |  |  |  |  |
| (a) | $\varphi = 0$ | $(\tau_Y)$ | $(\tau_R, \theta)$ | yes |
| (b) | $\beta = \frac{\tau_Y}{\tau_R} = 1$ | $(\varphi)$ | $(\tau_R, \theta)$ | yes |
| (c) | $\varphi = 1$ | $(\tau_R)$ | $(\tau_Y, \theta)$ | no |
| **(ii) Bidirectional introgression (BDI, Fig. 1D)** |  |  |  |  |
| (a) | $\varphi_X = 0, \varphi_Y = 0$ | $(\tau_Y)$ | $(\tau_R, \theta)$ | yes |
| (b) | $\varphi_X = 1, \varphi_Y = 1$ | $(\tau_Y)$ | $(\tau_R, \theta)$ | yes |
| (c) | $\beta = \frac{\tau_Y}{\tau_R} = 1$ | $(\varphi_X, \varphi_Y)$ | $(\tau_R, \theta)$ | yes |
| (d) | $\varphi_X = 0, \varphi_Y = 1$ | $(\tau_R)$ | $(\tau_Y, \theta)$ | no |
| (e) | $\varphi_X = 1, \varphi_Y = 0$ | $(\tau_R)$ | $(\tau_Y, \theta)$ | no |

Note.— For UDI, in segment c nuisance parameter $\tau_Y$ in $H_1$ is mapped onto $\tau$ in $H_0$ (Fig. 1A&B). For BDI, segments a, b, d, and e are lines, while segment c is a plane, and in segments d and e, nuisance parameter $\tau_Y$ in $H_1$ is mapped to $\tau_R$ in $H_0$ . In last column, ‘yes’ means that the condition on priors (eq. 5) holds for the segment and ‘no’ means it does not.

1. There exist three different routes via which *H*_1_ is reduced to *H*_0_ (by fixing the parameter of interest to its null value): (a) *φ* = 0, (b) *τ*_*Y*_ = *τ*_*R*_, and (c) *φ* = 1 (Fig. 1C).
2. For each route, certain parameters become unidentifiable or indeterminate when the parameter of interest takes the null value. For example, for route (a), *H*_1_ becomes *H*_0_ when *φ* = 0, but when *φ* = 0, the introgression time (*τ*_*Y*_) becomes unidentifiable; if there is no introgression, the time of introgression is irrelevant (Fig. 1C, table 1A).

Whereas in regular cases *H*_0_ corresponds to one point in the parameter space for *H*_1_, here *H*_0_ corresponds to three lines in *H*_1_ (Fig. 1D&E). Under such nonstandard conditions neither of the proofs of Dickey (1971) and Ji *et al*. (2023) works. Our first objective in this paper is thus to extend the theory for computing the Bayes factor via the S-D density ratio to nonstandard conditions.

Note that the irregularities discussed here affect the use of the S-D density ratio to calculate the Bayes factor, not the Bayes factor itself. However methods for calculating the Bayes factor via marginal likelihood (which are not affected by the irregularities discussed here) involve many MCMC runs and >100 times more computation than the S-D approach (Fourment *et al*., 2020, and see Discussion). We study simpler problems of Gaussian mixtures to develop the theory.

Our second objective is to study the behavior of the MCMC algorithm, influenced by the irregularities just mentioned. Figure 1F&G shows posterior summaries and trace plots for parameters in *H*_1_ from a typical bpp analysis of a large dataset simulated under the null model of no gene flow (*H*_0_, Fig. 1A). As the data are informative, the MCMC spends most of the time visiting the neighborhoods of the three lines of Figure 1D which represent the true model *H*_0_. However, do there exist multiple modes on the posterior surface (as the marginal summary of *φ* in Fig. 1F seems to suggest)? How often does the chain visit the neighborhoods of the three lines? How should one summarize the MCMC sample and what biological conclusions should one draw from such posterior samples? Our effort in this direction has not been completely successful, but we characterize the parameter space and posterior surface under *H*_1_, and the probabilities with which the MCMC visits different regions of the parameter space for *H*_1_ when large datasets are generated under *H*_0_ and analyzed under *H*_1_. We provide guidelines on Bayesian inference when gene flow between sister lineages is absent or occurs at very low rates. A recent simulation examined the robustness of the Bayesian test of gene flow between sister lineages to recombination and selection (Thawornwattana *et al*., 2026).

Our third objective is to evaluate the power of the Bayesian test and the precision of Bayesian estimation of the introgression probability, assessing the information content in multilocus sequence data concerning introgression between sister lineages.

Our paper is structured as follows. First we develop a theory for applying the S-D approach to computing the Bayes factor for testing introgression between sister lineages. We develop and illustrate the theory through simple examples, for which numerical calculation is efficient. We prove that when multiple routes exist for reducing *H*_1_ to *H*_0_, the local behavior of each route can be used to calculate the Bayes factor via the S-D approach. We study the posterior of parameters in the introgression model when large datasets are generated under the null model of no gene flow and suggest that the posterior probabilities for different regions of the parameter space may not converge to a point but rather to a non-degenerate probability distribution, when the data-size approaches infinity. We also highlight the extreme sensitivity of the posterior to the prior when there is no or little gene flow in the data. Finally we conduct a simulation study and an asymptotic analysis (treating data of infinite size) to examine the power of the Bayesian test of introgression and the precision of parameter estimation. We find that information content is greatly influenced by the species split time, with more information at deeper divergence. We discuss the impact of the irregularities of the testing problem on frequentist tests of introgression, such as the likelihood ratio test (LRT) based on the composite likelihood used in *∂*a*∂*i (Gutenkunst *et al*., 2009) and fastsimcoal2 (Excoffier *et al*., 2013).

## Theory

### Savage-Dickey (S-D) density ratio under nonstandard conditions

We consider the general case first and illustrate the theory using two simple examples involving Gaussian mixtures, before dealing with the Bayesian test of introgression between sister species.

Let the vector of parameters be ***θ***_0_ with the prior *π*_0_ (***θ***_0_) under *H*_0_, and ***θ*** with the prior *π* (***θ***) under *H*_1_. Given data ***x***, let the likelihood be *L*_0_ *θ*_0_ = *p*_0_ ***x*** (*θ*_0_ under *H*_0_ and *L* (***θ***) = *p* (***x*** | ***θ***) under *H*_1_. Here we use the subscript 0 for densities under *H*_0_ and no subscript for those under *H*_1_. The Bayes factor in support of *H*_1_ over *H*_0_ is defined as

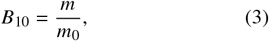

where *m* and *m*_0_ are the marginal likelihood values under the two models:

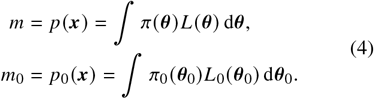

We recognize three types of parameters in *H*_1_, with ***θ*** = (*ω, λ, ξ*) , where *ω* are parameters of interest, which are constrained to formulate the null hypothesis (*H*_0_); *λ* are nuisance parameters, which are shared between *H*_0_ and *H*_1_ and usually have the same interpretation in the two models; and *ξ* are unidentifiable/indeterminate parameters, which become unidentifiable when *H*_1_ is reduced to *H*_0_.

In simple or regular cases, *H*_1_ reduces to *H*_0_ when the parameters of interest take special values that represent no effect (called *null value* or *null point*), that is, *ω* = *ω*_0_. For example, in the two-sample test (see SI text 1), samples are taken from two populations *N* (*μ*_1_, 1) and *N* (*μ*_2_, 1) to test the null hypotehesis *H*_0_ : *μ*_1_ = *μ*_2_ = *μ*. The parameter vector is ***θ***_0_ = (*μ*) in *H*_0_, and ***θ*** = (*μ*_1_, *μ*_2_) in *H*_1_, and *H*_1_ is reduced to *H*_0_ when *μ*_1_ = *μ*_2_ = *μ*. The parameter space for *H*_1_ is the 2-D *μ*_1_-*μ*_2_ plane, in which the diagonal line *μ*_1_ = *μ*_2_ represents the parameter space for *H*_0_ (which we refer to as the null space), while a point on the diagonal line (e.g., *μ*_1_ = *μ*_2_ = 0.5) is the null point or null value.

In contrast, in the test of introgression, there exist three routes from *H*_1_ to *H*_0_ (a, b, c in Fig. 1C), and the null point is not one point in the parameter space of *H*_1_ but rather corresponds to three lines or line segments, referred to later simply as segments (a, b, c in Fig. 1D&E). The null space (the parameter space for *H*_0_) corresponds to the three planes in Figure 1D&E.

Furthermore, the definition of the parameters of interest differ among the different segments, and as a result, the partitioning of the parameter space, ***θ*** = (*ω, λ, ξ*), differs depending on the route (table 1(i)).

We show that with multiple routes for reducing *H*_1_ to *H*_0_, each can be used to construct an S-D density ratio to compute *B*_10_.

#### Theorem 1.

*Suppose the null point (a point in the parameter space of H*_0_*) corresponds to multiple segments in the parameter space of H*_1_ *and at segment j* , *we partition the parameters as* ***θ*** = (*ω* _*j*_ , *λ* _*j*_ , *ξ* _*j*_), *where ω* _*j*_ *are parameters of interest (with ω* _*j*_ = *ω* _*j*0_ *being the null value), λ* _*j*_ *the nuisance parameters and ξ* _*j*_ *the unidentifiable parameters. On the condition that*

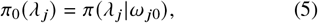

*where π* (*λ* _*j*_ | *ω* _*j*0_)= ∫*π* (*λ* _*j*_ , *ξ* _*j*_ | *ω* _*j*0_) d*ξ* _*j*_ , *the Bayes factor B*_10_ *of eq. 3 is given as*

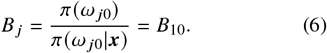

*Proof*. Note that *H*_0_ and *H*_1_ are nested, so that *L* (*ω* _*j*0_, *λ* _*j*_ , *ξ* _*j*_) = *L*_0_(*λ* _*j*_). Thus

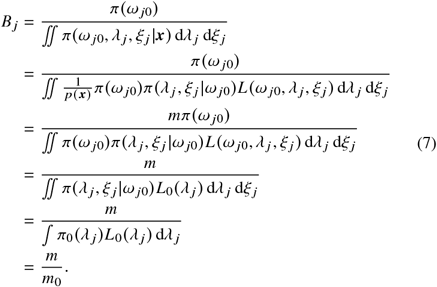

This proof resembles the reverse of the proof of Ji *et al*. (2023, eq. A1). However, here the argument concerns the local behavior of the densities around segment *j*. While the marginal likelihood values (*m*_0_, *m* in eq. 4) are defined by averaging over the whole parameter space in each model, the Bayes factor is given here by the local behavior of the posterior under *H*_1_ around the null point (or each of the segments that constitute the null point).

More generally, *H*_0_ may be specified by setting up equality constraints on parameters in *H*_1_, that is *c* (***θ***) = 0, rather than letting parameters of interest take the null value (*ω* = *ω*_0_). For example in the two-sample test we have ***θ*** = (*μ*_1_, *μ*_2_) , so the constraint is *c* (***θ***) = *μ*_1_ − *μ*_2_ = 0. As the parameters of interest depend on the segment, we use ***θ*** instead of *ω* _*j*_ to specify constraints. Suppose there are *J* routes of going from *H*_1_ to *H*_0_. For route *j* , let there be *K* _*j*_ equality constraints, 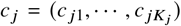, so that segment *j* is represented by

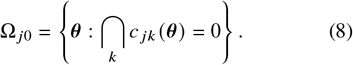

For simplicity, we use *K* instead of *K* _*j*_ but note that *K* may differ among segments (one such case is the test of bidirectional introgression; see table 1(ii)). We use infinitesimals *ϵ* _*jk*_ > 0, *j* = 1, …, *J*; *k* = 1, …, *K*, to define a *null region*, a subspace of the parameter space of *H*_1_ in the neighborhood of segment *j* ,

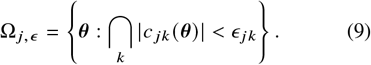

Let *ϵ* _*j*_ = (*ϵ* _*j*1_, …, *ϵ* _*jK*_). Define

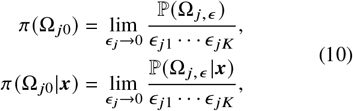

to be the prior and posterior densities for Ω _*j*0_ under *H*_1_. Then

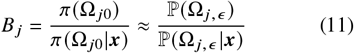

is an approximation of *B*_10_.

One may reparametrize *H*_1_ and treat *c* _*j*_ ≡ *ω* _*j*_ as parameters, and derive the induced prior on the new parameters. Then the Bayes factor for segment *j* of eq. 11 takes the simpler form of eq. 6. Thus we do not see any conceptual difference between eqs. 6 and 11, and will henceforth use the notation of eq. 11, possibly with reparametrization used to verify the condition on priors (eq. 5) (see Remark 2 below). Note that the number of constraints *K* in eqs. 10&11, which may be considered the degree of freedom for the test, equals the dimension of *ω* _*j*_ in eq. 6.

We will use the differentials *ϵ* _*j*_ to calculate P (Ω _*j,ϵ*_ | ***x***) for segment *j* by processing an MCMC sample under *H*_1_ (eq. 11) (Ji *et al*., 2023). One may also use kernel density smoothing to calculate *π* (Ω _*j*0_ | ***x***). The prior density *π* (Ω _*j*0_) is typically tractable analytically.

If the condition on priors holds for each segment, so that all segments give the same S-D density ratio, with *B* _*j*_ = *B*_10_ (eqs. 6 or 11), we can use all segments to calculate a combined density ratio. Let

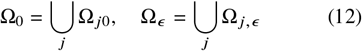

be the *null point* (or constraints) and *null region*, respectively. Then

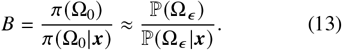

We make two observations regarding the condition on priors of eq. 5.

First, if the condition on priors (eq. 5) does not hold, a correction factor may be applied

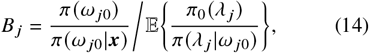

with the expectation taken over *π* (*λ* _*j*_ | ***x***, *ω* _*j*0_) (Verdinelli and Wasserman, 1995). This does not appear simpler to compute than the marginal likelihood values (*m*_0_, *m*), and is thus not pursued here. Instead later we discuss ideas for approximate calculation of *B*_10_ when the condition on priors does not hold.

Second, we note that the condition on priors (eq. 5) may hold for one parametrization but not another. Consider for example the one-sample test, in which *H*_0_ : *N* (0, *σ*^2^) is tested against *H*_1_ : *N μ, σ*^2^ with unknown variance *σ*^2^. The parameter of interest is *μ* with the null value *μ*_0_ = 0 while *σ*^2^ is the nuisance parameter. Conditioning on *μ* = 0 or on 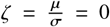 leads to different distributions: *p* (*σ*^2^|*μ* = 0) ≠ *p* (*σ*^2^|*ζ* = 0) (Fig. S1 and SI text 2). This phenomenon is known as the Borel-Kolmogorov paradox, in which conditioning on seemingly equivalent events leads to different conditional distributions. It is prudent to verify the condition on priors when calculating the S-D density ratio.

Note that the issue concerns the S-D approach to calculating the Bayes factor, not the Bayes factor itself. The marginal likelihood values are not affected by reparametrization, so that *B*_10_ is well-defined and is the same for different parametrizations.

### Example GM1. Gaussian mixture model for testing contamination

#### Problem setup

Here we consider a test of contamination under a Gaussian mixture model. This is not tractable analytically but yields to numerical calculation. It has most of the irregularities discussed above for the test of introgression, including multiple routes from *H*_1_ to *H*_0_, so we use it to illustrate the application of theorem 1 to calculate *B*_10_. We also use this example to study the posterior surface under *H*_1_ under nonstandard conditions.

The data consist of an i.i.d. sample, ***x*** = (*x*_*i*_), *i* =1, …, *n*. The likelihood model specifies *x*_*i*_ ∼ *N* (0, 1) under *H*_0_, or *x*_*i*_ ∼ *αN* (*μ*, 1) + (1 − *α*) *N* (0, 1) under *H*_1_, with probability *α* for contamination from *N* (*μ*, 1). Thus *H*_1_ is a mixture of two Gaussian distributions. We assign the prior *p α, μ* = *ϕ μ*; 0, 1 under *H*_1_, where *ϕ* (*z*; *μ, σ*^2^) is the probability density function (PDF) for the Gaussian distribution *N* (*μ, σ*^2^). There are no nuisance parameters, so the condition on priors (eq. 5) holds.

The marginal likelihoods under the two models are 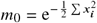

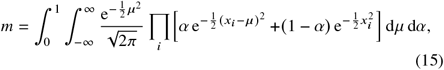

so that

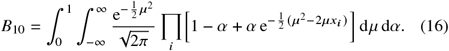

We use *m*_0_ to scale the likelihood under *H*_1_, and recognise the integrand in eq. 16,

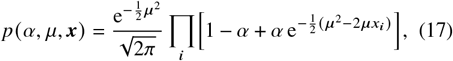

as the unnormalized posterior. Then the marginal likelihood for *H*_1_,

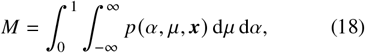

is also the Bayes factor: 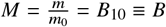.

#### The S-D approach to calculating B_10_

The constraint that reduces *H*_1_ to *H*_0_ is

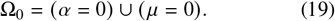

The null point comprises the two lines or axes of Figure 2A: (a) *α* = 0 (with *μ* unidentifiable) and (b) *μ* = 0 (with *α* unidentifiable). Define the null region

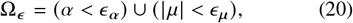

which are two narrow strips along the axes, of widths *ϵ* _*α*_ and 2*ϵ*_*μ*_, respectively (Fig. 2A). Each strip can be used to calculate the Bayes factor, giving

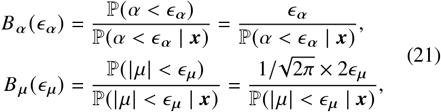

where the prior probabilities in the numerator are given by the uniform and Gaussian priors, while the posterior probabilities in the denominator can be estimated using an MCMC sample or calculated numerically using Gaussian quadrature.

**Fig. 2:**
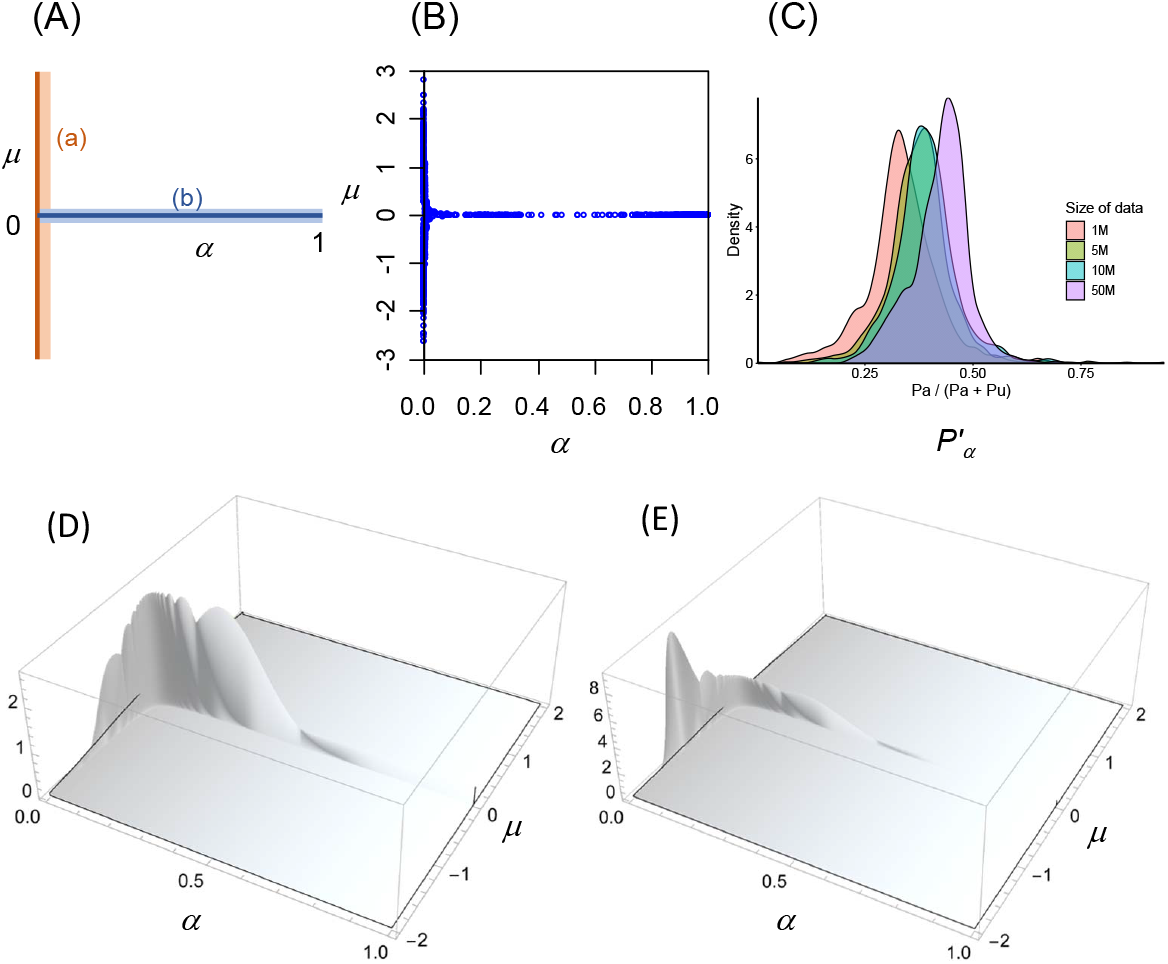
(**A**) Parameter space for the Gaussian mixture model *H*_1_ : *αN* (*μ*, 1) + (1 − *α*) *N* 0, 1) is the *α*–*μ* plane, with 0 ≤ *α* ≤ 1 and −∞ < *μ* < ∞. There are no unknown parameters in *H*_0_ : *N* (0, 1) , so that the null space is the null point and is represented by the two axes: (a) *α* = 0 (with *μ* unidentifiable) and (b) *μ* = 0 (with *α* unidentifiable). (**B**) MLEs 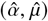 under *H*_1_ from 10^3^ datasets (of size *n* = 10^6^) simulated under *H*_0_. The MLE corresponds to the highest ℓ in ten runs of the BFGS algorithm. Note that all points on the two axes have identical likelihood so that the MLE is one unique point only if it is not on the axes. (**C**) Density for posterior probability for segment a (the *μ*-axis) at *n* = 10^6^, 5 × 10^6^, 10^7^, and 5 × 10^7^, estimated by simulating 1000 datasets under *H*_0_. Gaussian quadrature is used to calculate *P*_*α*_ and *P*_*μ*_ with 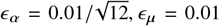, which are rescaled to give 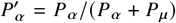. The average of *P*_*α*_ + *P*_*μ*_ is 0.5491, 0.6849, 0.7348, 0.8340, at the four values of *n*. Mathematica gave identical results to Gaussian quadrature for *n* ≤ 2000 but encounters numerical problems for larger *n*. (**D, E**) Unnormalized posterior surface (eq. 17) for two datasets of size *n* = 10^6^ simulated under *H*_0_. In the first (**D**), the sample mean is 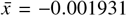, and the MLE under *H*_1_ is 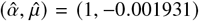 with the log likelihood (relative to that for *H*_0_) Δℓ = 1.864. Multiple local peaks exist close to the *α*-axis, including (0, any) with Δℓ = 0, and (0.328984, −0.005867) and (0.791298, −0.002440) both with Δℓ = 1.864. In the second (**E**), the sample mean is 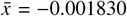. The MLE is (0.000927, −1.571305) close to the *μ*-axis, with Δℓ = 4.381, but local peaks exist close to the *α*-axis, such as (0.717088, −0.002553), (0.802798, −0.002280), and (0.909107, −0.002013) all with Δℓ = 1.675.

We also use both segments to produce a combined estimate, *B*_10_ ≈ ℙ (Ω_*ϵ*_)/ℙ (Ω_*ϵ*_ | ***x***).

#### The shape of the posterior surface

Next we study the posterior for parameters ***θ*** in *H*_1_ when large datasets generated under *H*_0_ are analyzed under *H*_1_. In a regular case, the posterior is expected to be concentrated around ***θ*** = ***θ***_0_ (the null value or the truth), with ***θ*** |***x*** ∼ *N* (***θ***_0_, (*nI*)^−1^), where ℐ is the Fisher information matrix, when *n* → ∞ (O’Hagan and Forster, 2004, p.72-74). Here we expect the posterior to be concentrated around the two segments (the *α*- and *μ*-axes) which constitute the null value. We are interested in the posterior probabilities for the two subregions around the two axes (Fig. 2A). They represent the proportional contributions of the two subregions to the marginal likelihood (*m*), and are also the proportions of times that the MCMC algorithm visits those subregions.

Let *ϵ* _*α*_ > 0 and *ϵ*_*μ*_ > 0 be fixed small values. We calculate the posterior probabilities for the two subregions *P*_*α*_ = ℙ (*α* < *ϵ* _*α*_ | ***x***) and *P*_*μ*_ = ℙ (|*μ*| < *ϵ*_*μ*_ ***x***|) using Gaussian quadrature. When *n* → ∞, both *P*_*α*_ and *P*_*μ*_ will increase, with *P*_*α*_ + *P*_*μ*_ → 1. We suggest that as *n* → ∞, *P*_*α*_ may not converge to a point value but instead vary among datasets (all generated under *H*_0_) according to a non-degenerate distribution. Here we provide without proof an intuitive argument through analogy and generate the distribution of *P*_*α*_ numerically.

The problem appears to be similar to the fair-coin or fair-balance paradox, the star-tree paradox (Lewis *et al*., 2005; Yang and Rannala, 2005; Yang, 2007) and in general comparison of equally correct and nondistinct models (Yang and Zhu, 2018, Fig. B2). In the fair-balance paradox, a balance is fair but we test whether it has a negative or positive bias (Yang and Rannala, 2005). We compare two models, *H*_−_ : *μ* < 0 and *H*_+_ : *μ* ≥ 0 when the truth is *μ* = *μ*_0_ = 0. The data are an i.i.d. sample of size *n*, generated from *x*_*i*_ ∼ *N* (0, 1) and summarized as the sample mean 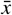. With the prior *μ* ∼ *N* (0, 1), the posterior model probability is

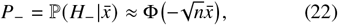

where Φ is the cumulative distribution function (CDF) for *N* (0, 1) (Yang and Rannala, 2005, eq. 6). As 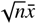 is an *N* (0, 1) variable, *P*_−_ has a *U*(0, 1) distribution.

As *n* is large, 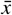 is very close to *μ*_0_ = 0 in essentially every dataset. However a small deviation of 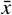 from 0 can translate to a large departure of *P*_−_ from 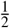. Let *n* = 10^6^ , say. Then in 2.5% of datasets 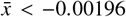 so that *P*_−_ > 0.975, while in another 2.5% of datasets 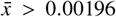 so that *P*_−_ < 0.025. Indeed as 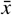 varies among datasets according to 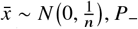 varies like a random number from *U*(0, 1).

Note that the Bayesian test of *H*_0_ : *N* (0, 1) against *H*_1_ : *N* (*μ*, 1), with the prior *μ* ∼ *N* (0, 1), produces

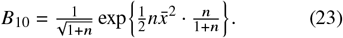

As 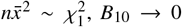 when *n* → ∞. For example, at *n* = 10^6^ and 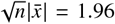, we have *B*_10_ = 0.00682, strongly supporting *H*_0_. However, 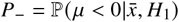 fluctuates among datasets like a random number.

In the Gaussian-mixture model of Figure 2, when *n* is large enough the posterior probability outside the two strips (represented by Ω_*ϵ*_) will be ∼ 0 with *P*_*α*_ + *P*_*μ*_ ≈ 1. The two strips are like the parameter space for the two models *H*_−_ and *H*_+_ in the fair-balance example, with MLEs under *H*_1_ close to the *α* and *μ* axes corresponding to 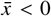 or 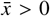. Thus the posterior surface under *H*_1_ may be expected to vary among replicate datasets (all simulated under *H*_0_) in terms of the probability mass above the two strips. Figure 2D&E shows the unnormalized posterior for two datasets, each of size *n* = 10^6^ simulated under *H*_0_. In the first, the MLE is close to the *α*-axis, while in the second, the MLE is close to the *μ*-axis. Note that in all large datasets, the MLEs are close to the two axes, supporting the null model of no contamination (Fig. 2B).

We simulated 1000 datasets for a few large *n* values (from 10^6^ to 5 × 10^7^) under *H*_0_ : *N* (0, 1) , and calculated *P*_*α*_, *P*_*μ*_, with the histogram for the rescaled probability, 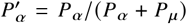, shown in Figure 2C. Even with very large 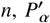 varies among datasets according to a distribution rather than converging to a point value.

### Example GM2. Another Gaussian-mixture model

*Problem setup*. We modify the previous example slightly, so that *H*_0_ specifies one population with mean *μ* while *H*_1_ specifies two populations with means *μ*_1_ and *μ*_2_ in proportions *α* and 1 − *α* (Fig. 3A). This example has all the irregularities encountered in the test of unidirectional introgression of Figure 1A&B and indeed the two problems have the same geometrical structure (compare Fig. 3A with Fig. 1D). This example is not analytically tractable but the MCMC algorithm involves much less computation than the test of introgression (compare Fig. 3B–D with Fig. 1F&G).

**Fig. 3:**
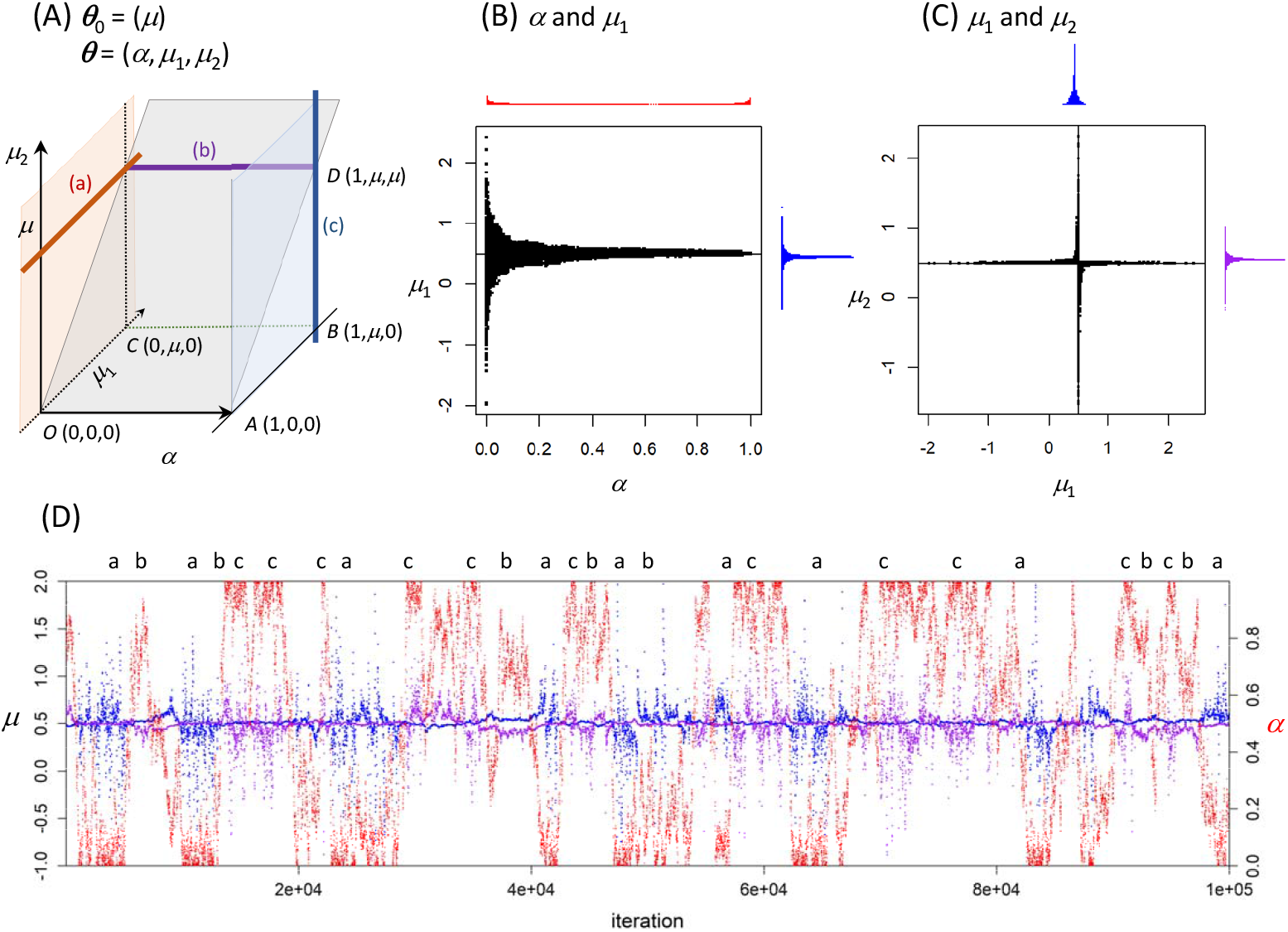
(**A**) The 3-D parame→ter space for the Gaussian mixture model *H*_1_ : *αN* (*μ*_1_, 1) + (1 − *α*) *N* (*μ*_2_, 1), with parameter vector ***θ*** = (*α, μ*_1_, *μ*_2)_. The null model is *H*_0_ : *N* (*μ*, 1) , with ***θ***_0_ = (*μ*). The parameter space of *H*_0_ (the null space) corresponds to the three planes while one point in the parameter space of *H*_0_ (the null point or a particular *μ* value such as *μ* = 0.5) corresponds to three lines or line segments: (a) *α* = 0 (with *μ*_1_ unidentifiable and with *μ*_2_ → *μ*), (b) *μ*_1_ = *μ*_2_ (with 0 ≤ *α* ≤ 1 unidentifiable and with *μ*_1_ = *μ*_2_ → *μ*), and (c) *α* = 1 (with *μ*_2_ unidentifiable and with *μ*_1_ → *μ*). (**B, C**) 2-D scatter plots and (**D**) trace plots of *α* (red), *μ*_1_ (blue) and *μ*_2_ (purple) from an MCMC analysis under *H*_1_ of a dataset of size *n* = 10^5^, simulated under *H*_0_ with *μ* = 0.5. Labels a, b, c on top of the trace plot (**D**) indicates that the chain is around the three segments of panel **A**.

The parameter vector is ***θ***_0_ = *μ* for *H*_0_ and ***θ*** = (*α, μ*_1_, *μ*_2)_ for *H*_1_ (Fig. 3A). This is again a test for mixture. The data are an i.i.d. sample, ***x*** = (*x*_*i*_), with *x*_*i*_ ∼ *N* (*μ*, 1) under *H*_0_, or *x*_*i*_ ∼ *αN* (*μ*_1_, 1) + (1 − *α*) *N* (*μ*_2_, 1) under *H*. The prior under *H* is 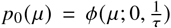, with *τ* = 1. Under *H*_1_ we assign the prior *U*(0, 1) for *α* and 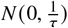 for *μ*_1_ and *μ*_2_, so that

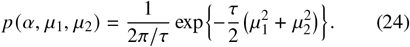

If we re-parametrize *H*_1_ and use ***θ***^′^ = (*α, μ*_1_, *δ*) instead, with *δ* = *μ*_2_ − *μ*_1_, the prior becomes 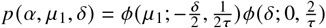.

Under *H*_1_, the density for *x* is

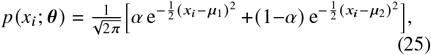

so that the posterior is

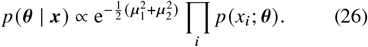

There are three routes from *H*_1_ to *H*_0_, and the constraints that reduce *H*_1_ to *H*_0_ are

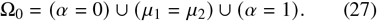

The null point corresponds to three lines or line segments in the parameter space of *H*_1_: (a) *α* = 0 (with *μ*_1_ unidentifiable), (b) *μ*_1_ = *μ*_2_ (with *α* unidentifiable), and (c) *α* = 1 (with *μ*_2_ unidentifiable). The parameter space for *H*_0_ (the null space) corresponds to three planes in the parameter space of *H*_1_ (Fig. 3A).

We note that the condition on priors (eq. 5) holds for segments (a) *α* = 0 and (c) *α* = 1, but not for segment (b) *μ*_1_ = *μ*_2_ or *δ* = *μ*_2_ − *μ*_1_ = 0. For segment (b), we have 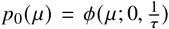 for *H*_0_, and 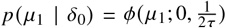 for*H*,sothereisamis-match.

#### Bayes factor via the S-D density ratio

We use the differentials *ϵ* _*α*_ and 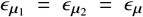 to define a null region in the parameter space of *H*_1_,

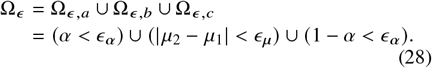

If we use each segment to calculate *B*_10_, we have

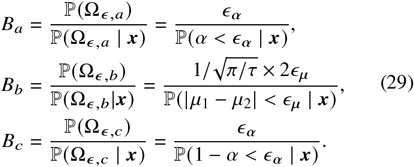

We use the MCMC sample of Figure 3D to calculate the S-D density ratio with different differentials *ϵ* _*α*_ and *ϵ*_*μ*_ (Fig. S2B). We expect *B*_*a*_ and *B*_*c*_ to approximate *B*_10_ but *B*_*b*_ may be off by a factor, since the condition on priors holds for segments a and c but not for b. However, for this dataset the three segments produced similar *B*_10_ values, around 0.25, with the data preferring the null model of no mixture (i.e., the true model).

If we use all segments to calculate a combined S-D density ratio, the prior probability for the null region under *H*_1_ will be

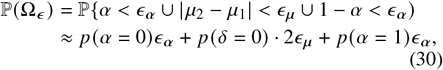

while the posterior probability ℙ (Ω_*ϵ*_ | ***x***) can be calculated by processing the MCMC sample under *H*_1_. As the different segments produced similar *B*_10_ estimates in the example of Figure 3D (Fig. S2B), the combined estimate is similar as well.

#### The shape of the posterior surface

Under *H*_1_, ***θ*** = (*α, μ*_1_, *μ*_2_) and ***θ***^′^ = (1 − *α, μ*_2_, *μ*_1_) are unidentifiable and this is an unidentifiability of the label-switching type. As we use a symmetrical prior *α* ∼ *U* (0, 1) , the posterior *p* (*α* | ***x***) around segments a and c are mirror images of each other. We thus consider the relative posterior probabilities around segments a and c versus segment b (Fig. 3A). Rousseau and Mengersen (2011) analyzed the posterior under an over-fitting finite-mixture model with too many components when large datasets are generated under the null model and when the mixing probabilities are assigned a Dirichlet prior. Example GM2 is case 1 analyzed in Rousseau and Mengersen (2011, Fig. 1). The dynamics is found to depend on the parameters of the Dirichlet prior in relation to the number of parameters in each component (*d*). If *a* < *d*/2, the posterior probability that the extra component has non-negligible probability (averaged over datasets) is small, or in other words the posterior will be dominated by ‘emptying’ a component (with either *α* or 1 − *α* being ≈ 0, i.e., segments a or c in Fig. 3A). If *a* > *d*/2 the posterior will be dominated by ‘merging’ components (with *μ*_1_ = *μ*_2_ or segment b in Fig. 3A). The analysis highlights the influence of the prior on the mixing probabilities on the posterior. In our example *a* = 1 and there is one parameter in each component (e.g., *μ*_1_ for the first component), so that *a* > *d*/2. The authors simulated small datasets only with *n* ≤ 10^3^. Our simulation of larger datasets (*n* = 10^6^) shows a U-shaped posterior for *α*, and segment b does not dominate the posterior. We leave it to future research to investigate the impact of the data and prior on the posterior when data are generated under *H*_0_ and analyzed under *H*_1_.

### Test of unidirectional introgression (UDI) between sister lineages

The test of introgression (under the UDI model) has been introduced in Introduction (Fig. 1; see also Fig. 4A&B). The null model *H*_0_ assumes no gene flow, with the parameter vector ***θ***_0_ = (*τ, θ*), while the alternative model *H*_1_ assumes unidirectional introgression, with the parameter vector ***θ*** = (*φ, τ*_*Y*_ , *τ*_*R*_, *θ*) (Fig. 1B). Here we assume the same population size *θ* for all species on the phylogeny, which has the same prior under *H*_0_ and *H*_1_. In models with different population sizes, the *θ* parameters have the same priors in the two models as well. There are three routes from *H*_1_ to *H*_0_, and the null point (one point in the parameter space of *H*_0_) corresponds to three lines or line segments in the parameter space of *H*_1_ given by the constraints

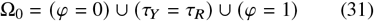

(table 1(i), Fig. 1D). The null space (the parameter space for *H*_0_) is represented by the three planes in Figure 1D.

**Fig. 4:**
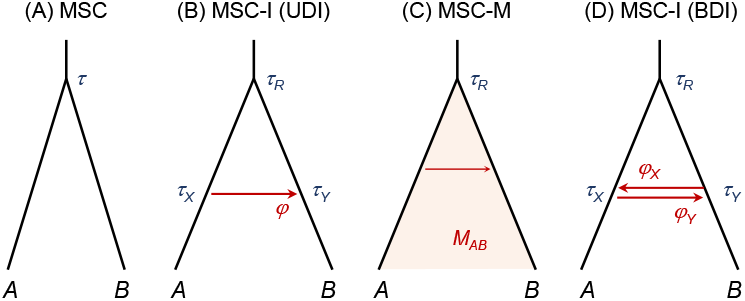
(**A**) Unidirectional introgression model (MSC-I, UDI) with *A* → *B* (or *X* → *Y*) introgression occurring at time *τ*_*X*_ ≡ *τ*_*Y*_ and introgression probability *φ*_*Y*_. (**B**) Continuous migration model (MSC-M) with *A B* migration at the population migration rate *M*_*AB*_ per generation. (**C**) Bidirectional introgression model (MSC-I, BDI) with introgression probabilities *φ*_*X*_ and *φ*_*Y*_ and introgression time *τ*_*X*_ ≡ *τ*_*Y*_. The MSC-I UDI and MSC-M models (**A, B**) are used to simulate data while all three models of gene flow (**B**-**D**) are used to analyze data.

In bpp, we assign a beta prior *φ* ∼ Beta (*a, b*) , a gamma-Dirichlet prior on the split times (Yang and Rannala, 2010, eq. 2), which means that *τ*_*R*_ ∼ *G* (*α*_*τ*_, *β*_*τ*_), and given *τ*_*R*_, *β* = *τ*_*Y*_/*τ*_*R*_ is *U* (0, 1). We note that the condition on priors (eq. 5) holds for segments a and b, but not for segment c (table 1(i)). For segments a and b, the nuisance parameters are *τ*_*R*_ and *θ* , each with the same gamma prior in both *H*_0_ and *H*_1_. For segment c, the nuisance parameter *τ*_*Y*_ in *H*_1_ is mapped onto *τ* in *H*_0_, with different priors. In *H*_0_ the prior is *τ* ∼ *G*(*α, β*), with the prior mean *α*/*β*. In *H*_1_, we have *τ*_*Y*_ |*τ*_*R*_ ∼ *U*(0, 1) with *τ*_*R*_ ∼ *G*(*α, β*) so that

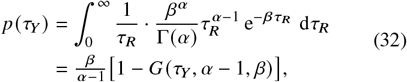

where *G* (*y, α, β*) is the CDF for the gamma distribution *G* (*α, β*) (Flouri *et al*., 2022). This has the prior mean *α*/(2*β*).

From eq. 31, we define the null region

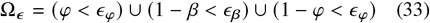

to calculate the Bayes factor, 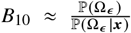, where ℙ (Ω_*ϵ*_) = *ϵ*_*φ*_ + *ϵ*_*β*_ + *ϵ*_*φ*_ while ℙ (Ω_*ϵ*_ |***x***) is the proportion of MCMC samples under the UDI model that fall in the null region (Ω_*ϵ*_).

We use this approach in our simulation below to evaluate the performance of the Bayesian test, setting the differentials to be 0.01*σ* with *σ* to be the prior standard deviation; in other words we use 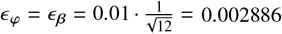.

#### The shape of the posterior surface and the impact of the prior

As mentioned in the Introduction, when large datasets generated under the null model of no gene flow (*H*_0_, Fig. 1A) are analyzed under the UDI model (*H*_1_, Fig. 1B), the posterior tends to be concentrated around the three segments (a, b, c in Fig. 1D&E) that collectively represent the true model of no gene flow. Qualitatively, the asymptotics under the UDI model appears to be similar to that under the mixture models (GM1 and GM2), but it is unclear whether the asymptotic analysis of finite-mixture models by Rousseau and Mengersen (2011) applies to the introgression model. In the special case of sampling one sequence from the recipient species, the UDI model specifies a mixture of gene trees and the likelihood model is a mixture model with mixing probabilities *φ* and 1 − *φ* for two components. When two or more sequences are sampled from the recipient species the data are not described by a mixture model anymore (see Meng and Kubatko, 2009 and correction in Jiao *et al*., 2021).

Similarly to Rousseau and Mengersen (2011), we have observed high sensitivity of the posterior to the prior on the introgression probability (*φ*) under the UDI model when data are generated with no gene flow. We analyzed the data of Figure 1 under the UDI model using the prior *φ* ∼ Beta(*a, a*) with 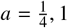, and 10 (Fig. S3). The Beta(*a, a*) prior has a U-shape when *a* < 1, is flat when *a* = 1, and has a bell shape when *a* > 1. At 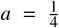 or 1, the posterior has substantial mass close to segments a and c, with *φ* close to 0 or 1. At *a* = 10, the posterior shifts to segment b, away from 0 and 1. In Bayesian analyses under regular conditions, the prior typically becomes less and less important when the amount of data increases, and eventually the posterior is entirely dominated by the data when *n* → ∞. Here the prior has an effect on the posterior even when *n* → ∞ (Rousseau and Mengersen, 2011). Nevertheless, it should be noted that the impact of the prior is to shift the posterior mass among three segments of Figure 1D&E, all of which represent the same likelihood model and the same biological scenario of no gene flow.

### Test of bidirectional introgression (BDI) between sister lineages

Under the BDI model there are two introgression probabilities, so that the parameter vector in *H*_1_ is ***θ*** = (*φ*_*X*_, *φ*_*Y*_ , *τ*_*Y*_ , *τ*_*R*_, *θ*) (Fig. 4D). The null model *H*_0_ is the MSC model with no gene flow as before, with ***θ***_0_ = (*τ, θ*) for *H*_0_. The common population size parameter *θ* is a nuisance parameter that exists in both models, with the same prior.

There are five routes by which *H*_1_ can be reduced to *H*_0_. In other words, one point in *H*_0_ (the null point) corresponds to five lines (or plane) specified by equality constraints on four parameters in *H*_1_: *φ*_*X*_, *φ*_*Y*_ , *τ*_*Y*_ , and *τ*_*R*_ (table 1(ii)),

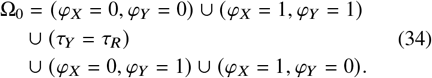

The condition on priors (eq. 5) holds for segments a–c but not for segments d and e (table 1(ii)). For the latter two, the nuisance parameter *τ*_*Y*_ in *H*_1_, which has the prior of eq. 32, is mapped onto *τ* in *H*_0_, with the prior *τ* ∼ *G*(*α, β*), so there is a mismatch.

From eq. 34, define the null region as

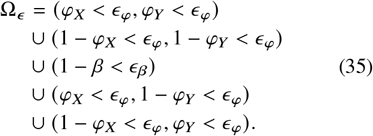

As in the test of UDI, we set *ϵ*_*φ*_ = *ϵ*_*β*_ = 0.002886 to process the MCMC sample from the BDI model to calculate ℙ (Ω_*ϵ*_ |***x***). For the prior probability, we have

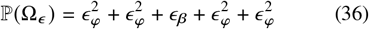

We use this approach in the computer simulation below.

### Test of gene flow between sister lineages under continuous migration model (MSC-M)

We consider two variants of the continuous MSC-migration (MSC-M) model of gene flow between sisters: the isolation-with-migration (IM) model in which migration has been ongoing until the present time (Fig. 4C) and the isolation-with-initial-migration (IIM) model (Costa and Wilkinson-Herbots, 2017; see also Huang *et al*., 2022, Fig. 1B), which assumes that gene flow occurs post speciation initially but ceased at a certain time in the past (*τ*_*Y*_ , Fig. 5B&C). The Bayesian test of gene flow under the IM model does not involve the irregularities discussed in this paper but the IIM model involves similar irregularities (table 2).

**Table 2:**
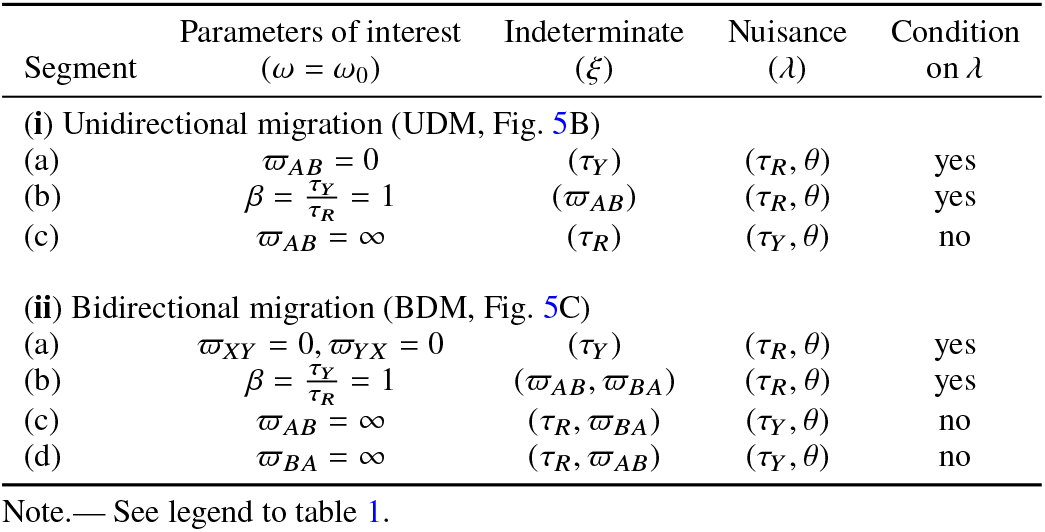
Parameters of interest (*ω*), nuisance parameters (*λ*), and indeterminate parameters (*ξ*) and condition on priors in the Bayesian test of gene flow under the isolation-with-initial-migration (IIM) models

**Fig. 5:**
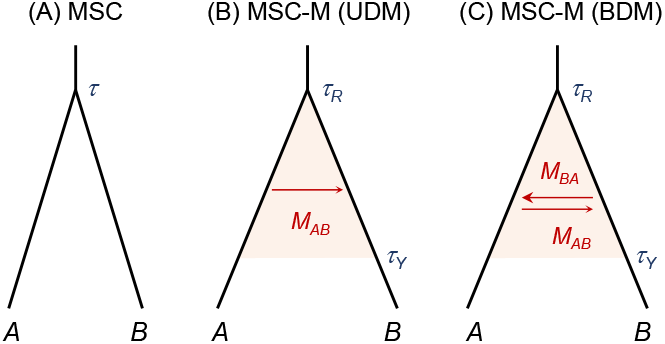
(**A**) The null model of no gene flow (*H*_0_: MSC). (**B, C**) Unidirectional and bidirectional migration (UDM, BDM) models with gene flow stopping at time *τ*_*Y*_. Both instances of the MSC-M model assumes isolation with initial migration (IIM) (Costa and Wilkinson-Herbots, 2017; see also Huang *et al*., 2022, Fig. 1B).

First we note that version 4.8 of BPP introduced a re-parametrization so that the mutation-scaled migration rate *ϖ*_*XY*_ = *m*_*XY*_/*μ* = 4*M*_*XY*_/*θ*_*Y*_ is used instead of the population migration rate *M*_*XY*_ = *m*_*XY*_ *N*_*Y*_ , where *m*_*XY*_ is the expected proportion of *X* → *Y* immigrants in the recipient population *Y* , so that *M*_*XY*_ and *ϖ*_*XY*_ are the expected number of *X* → *Y* immigrants per generation and per mutational time unit, respectively. Gamma priors are assigned to *ϖ* while *θ*s are assigned independent gamma or inverse-gamma priors.

Under the IM model, *H*_0_ is represented by *ϖ*_*AB*_ = 0 in *H*_1_. This null value is at the boundary of the parameter space (which is not a problem for either the Bayes factor or the S-D approach for calculating it), but no other parameters become indeterminate. There do not exist other routes from *H*_1_ to *H*_0_. For example, letting *φ* = reduces *H*_1_ to the model of one species (or two species with *τ*_*r*_ = 0) but not *H*_0_, which is the complete-isolation model for two species (Fig. 5A). The bidirectional migration model in the case of two species (BDM, with both *ϖ*_*AB*_ and *ϖ*_*BA*_) is also identifiable with data of multiple samples per species per locus, unlike the discrete introgression model (BDI), which shows two unidentifiable modes (Yang and Flouri, 2022).

Under the IIM model, if gene flow is unidirectional (UDM), the situation for the test is very similar to that under the discrete UDI model, and there exist three routes by which *H*_1_ is reduced to *H*_0_: (a) *ϖ* = 0 (with *τ*_*Y*_ unidentifiable), (b) *τ*_*Y*_ = *τ*_*R*_ (with *ϖ* unidentifiable), and (c) *ϖ* = 1 (with *τ*_*R*_ unidentifiable) (Fig. 5A&B, table 2(i)). Similarly to the test under UDI (Fig. 1), we can calculate the Bayes factor using the Savage-Dickey density ratio based on each of the three segments. For example, for segment a, 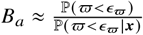, with *ϵ* _*ϖ*_ to be 0.5% or 0.1 % quantile for the gamma prior *ϖ* ∼ *G*(*α, β*). Segment c may be used to calculate 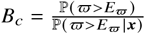, where *E* _*ϖ*_ may be set to the 99.5% or 99.9% quantile for the gamma prior.

Under the IIM model with bidirectional gene flow (BDM), there exist four routes by which *H*_1_ is reduced to *H*_0_ (Fig. 5A&C, table 2(ii)). Segments (a) and (b) are the same as in the case of UDM, but there are two other routes (segments c and d). If any of the two migration rates (*ϖ*_*AB*_, *ϖ*_*BA*_) is , *H*_1_ will become *H*_0_, with *τ*_*Y*_ ∞ → *τ* and with the other migration rate becoming indeterminate (as is *τ*_*R*_). We leave it to the future to explore the posterior surface when large datasets of no gene flow are analyzed under the IIM model.

## Materials and Methods

### Simulation under the null model of no gene flow

To verify our theory for applying the S-D approach to calculating the Bayes factor and to examine the false-positive rate of the Bayesian test of gene flow, we conducted a small simulation under the null model of MSC with no gene flow (Fig. 1A or Fig. 4A). We used the population size *θ* = 0.005 and three values for the species split time: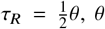, and 10*θ*, representing very shallow, shallow, and deep divergences, respectively. Note that 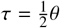 represents a very shallow phylogeny as 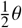 is the average coalescent time between two sequences from the same species. Each dataset consisted of *L* = 500 loci, with 10 haploid sequences per locus per species, and with the sequence length to be 500. The number of replicates was 100. The `simulate` option of bpp was used to generate gene trees for the loci and then to ‘evolve’ sequences along branches of the gene trees under the JC model (Jukes and Cantor, 1969) to generate sequence alignments.

Each dataset was analyzed using bpp under the MSC-I (UDI and DBI) and MSC-M models of Figure 4B–D to conduct the Bayesian test. In analyses under the MSC-I models, the age of the root *τ*_*R*_ was assigned a gamma prior *G* (2, 400) , with the prior mean 0.005, too large for data simulated using 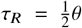 and too small for *τ*_*R*_ = 10*θ*. Given *τ*_*R*_, *τ*_*X*_ under UDI and BDI has the uniform distribution. The option `thetamodel = linked-all` in bpp was used so that the same population size (*θ*) was assumed for all populations on the species tree, and assigned the gamma prior *θ* ∼ (*G* 2, 400) with mean 0.005. Note that models with different *θ*s for species on the phylogeny are available in bpp, but in this paper assumptions on *θ*s are unimportant. The introgression probability *φ* (or both *φ*_*Y*_ and *φ*_*X*_ in the BDI model) was assigned the prior *U* (0, 1).

We used a burn-in of 40,000 iterations, after which we took 10^5^ samples, sampling every 2 iterations. Convergence is assessed by running the same analysis twice and confirming consistency between runs. Running time for analyzing one dataset was ~ 11 hours using one thread.

The MCMC sample under *H*_1_: MSC-I was processed to calculate the Bayes factor via the S-D density ratio. For the test of *H*_0_: MSC against *H*_1_: MSC-I (UDI), we used 1% of the prior standard deviations as the differentials, so that 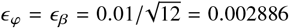. From eq. 31, the null region is

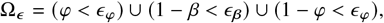

where *β* = *τ*_*Y*_/*τ*_*R*_ (eq. 31). We calculate the Bayes factor as 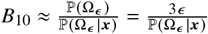.

We also tested the idea of using segments *a* and *b* only, for which the condition on priors holds (eq. 5), with

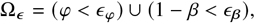

and 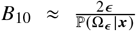. This produced nearly identical results as the use of all three segments.

For testing BDI, we used the null region defined in eq. 34. As in the test of UDI, we set *ϵ*_*φ*_ = *ϵ*_*β*_ = 0.002886. We have (see eq. 36)

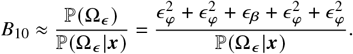

If we use only segments *a, b, c*, for which the condition on priors hold (table 1(i)), we have

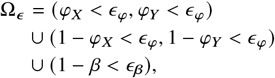

and 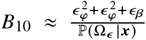. This produced nearly identical results as the use of all five segments.

The same data, simulated under *H*_0_, were also analyzed under the migration model (*H*_1_: MSC-M, Fig. 4C) to conduct the Bayesian test. The MSC-M model assumes that migration from *A* → *B* occurs continuously in every generation (Flouri *et al*., 2023). We used two priors to assess the impact of the prior, *ϖ*_*XY*_ ∼ *G* (2, 0.01) with mean 200 and *G* (2, 1) with mean 2. Like the analysis under the MSC-I models (UDI and BDI), we assumed the same population size for all species (`thetamodel = linked-all`). The S-D approach was used to calculate the Bayes factor *B*_10_, using the null region Ω_*ϵ*_ : *ϖ* < *ϵ* _*ϖ*_ = 0.005*σ*, 0.01*σ*, 0.05*σ*, where *σ* is the prior standard deviation. Then *B*_10_ ≈ P(Ω_*ϵ*_)/P(Ω_*ϵ*_ |***x***), with P(*ϖ* < *ϵ* _*ϖ*_) given by the CDF for the gamma prior.

### Simulation under models of gene flow

To assess the power of the Bayesian test of gene flow between sister lineages and the precision of Bayesian estimation of the rate of gene flow, we simulated data under the MSC-I and MSC-M models of Figure 4B&C, with *A* → *B* (or *X* → *Y*) gene flow. Most settings were the same as for the simulation under the null model. As before we used three species split times: 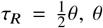, and 10*θ*, with *θ* = 0.005. Under the MSC-I model, the introgression time was set at 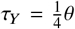, while the introgression probability varied: *φ* = 0.05, 0.1, 0.2, 0.3, 0.5, 0.7. For the MSC-M model, the population migration rate (from *A* → *B*) was *M* = 0.05, 0.1, 0.2, 0.3, 0.5, 1. Each dataset consists of *L* = 500 loci, with 10 + 10 sequences, with the sequence length 500. The number of replicates is 100.

Each replicate dataset was analyzed under the three models of gene flow of Figure 4B–D: MSC-I (UDI), MSC-I (BDI), and MSC-M. One population size (*θ*) was assumed for all species. The priors and MCMC settings were as above for analysis of data simulated under the null model. The Bayesian test of gene flow was conducted as before, using the S-D density ratio to calculate the Bayes factor.

### Simulation under the Gaussian mixture model (GM2)

A C program is written to generate MCMC samples under the Gaussian-mixture model *H*_1_ : *αN* (*μ*_1_, 1) + (1 − *α*) *N* (*μ*_2_, 1)to estimate parameters ***θ*** = (*α, μ*_1_, *μ*_2_) (Fig. 3**A**). A dataset of size *n* = 10^5^ was simulated under *H*_0_ : *N μ*, 1 with *μ* = 0.5. The MCMC algorithm includes sliding-window steps to update *α, μ*_1_, and *μ*_2_ separately, with another step updating *μ*_1_ and *μ*_2_ together. Window sizes are proportional to 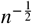. Besides acceptance rates of the moves, one can also use features of the posterior to assess convergence, as the posterior means 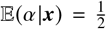, and 𝔼 (*μ*_1_|***x***) = 𝔼 (*μ*_2_|***x***). Running time for 10^6^ MCMC iterations was 1h20m using one thread on a server.

### Bayesian inference of gene flow between two lizard species

We analyzed a dataset of 1000 ddRAD loci from three Spiny lizard species from Baja California, *Sceloporus orcutti* (O), *S. hunsakeri* (H), and *S. licki* (L), published and analyzed by Gottscho *et al*. (2025). There are 16, 16, and 12 haploid sequences per locus for O, H, and L, respectively, with the sequence length to be ∼90bps. The species phylogeny is shown in Figure 6A, with possible gene flow between *S. orcutti* and *S. hunsakeri*.

**Fig. 6:**
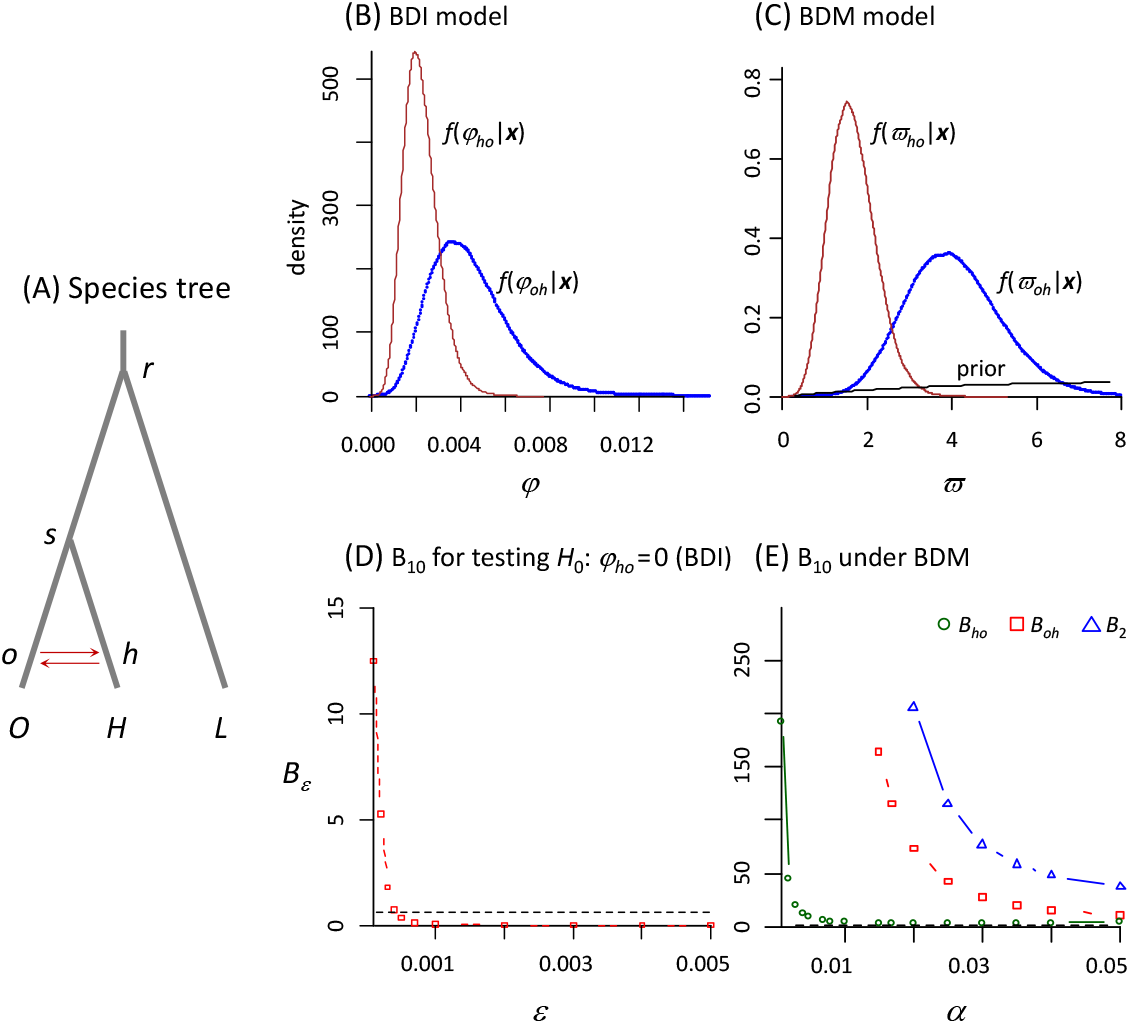
(**A**) Phylogeny for three Spiny lizard species, *Sceloporus orcutti* (O), *S. hunsakeri* (H), and *S. licki* (L), with gene flow between O and H. (**B**) Posterior densities for introgression probabilities *φ*_*oh*_ and *φ*_*ho*_ obtained in bpp analysis of the ddRAD data under the BDI model. The prior *φ* ∼ *U* 0, 1) is invisible. (**C**) Posterior densities for migration rates (*ϖ*_*oh*_, *ϖ*_*ho*_) under the BDM model. The prior *ϖ* ∼ *G* (2, 0.1) is shown as well. See table 5 for estimates of other parameters under the two models. (**D**) Approximate Bayes factor (*B*_10,*ϵ*_) for testing *H*_0_ : *φ*_*ho*_ = 0 against *H*_1_ : *φ*_*ho*_ ≠ 0, with *φ*_*oh*_ treated as a nuisance parameter, calculated using an MCMC sample under BDI of panel B. (**E**) *B*_10,*ϵ*_ for testing *H*_0_ : *ϖ*_*ho*_ = 0; *H*_0_ : *ϖ*_*oh*_ = 0; or *H*_0_ : *ϖ*_*ho*_ = 0, *ϖ*_*oh*_ = 0 using the MCMC sample under BDM of panel **D**, plotted against the tail probability of the prior, *α* = ℙ (*ϖ* < *ϵ*).

We assigned the gamma prior *τ*_*r*_ ∼ *G* (2, 200) with mean 0.01 on the age of the root, *θ* ∼ *G* (2, 400) with mean 0.005 for the population sizes, and the uniform prior *U* (0, 1) on the introgression probabilities (*φ*_*oh*_, *φ*_*ho*_). For the MCMC, we used a burn-in of 40,000 iterations, after which we took 2 × 10^5^ samples, sampling every 2 iterations. The run took 11hrs using 36 threads on a server. The MCMC samples were processed to calculate the Bayes factor for testing the hypothesis of no gene flow using the S-D density ratio, as described in the paper.

The same data were also analyzed under the MSC-M model (IM) with bidirectional migration. The mutation-scaled migration rates (*ϖ*_*oh*_, *ϖ*_*ho*_) were assigned the gamma prior *G* (2, 0.1) with mean 20. Other settings were the same as for the BDI model. Running time was 15hrs using 36 threads.

We also fitted the IIM model, specified as an instance of the MSC-M model by using a ghost lineage in the species tree for which no data are available, `((O, (H, ghost) h) s, L) r;` and by specifying migration between lineage *O* and *h* (the parent node of *H* and ghost) (Fig. 6**A**). The option `thetamodel = linked-mscm` was specified to force branches *Hh* and *hs* to share the same *θ*_*H*_. Other settings are the same as for the IM model.

## Results

### Bayesian test of gene flow between sister species has low false positives

Results for the Bayesian test of gene flow using data simulated under the null model (MSC, Fig. 4A) are summarized in table 3. We discuss the results with *ϵ*_*φ*_, *ϵ*_*β*_ in MSC-I and *ϵ* _*ϖ*_ in MSC-M set to 0.01*σ*, where *σ* is the prior standard deviation. Use of 0.005*σ* and 0.05*σ* produced very similar results.

**Table 3:**
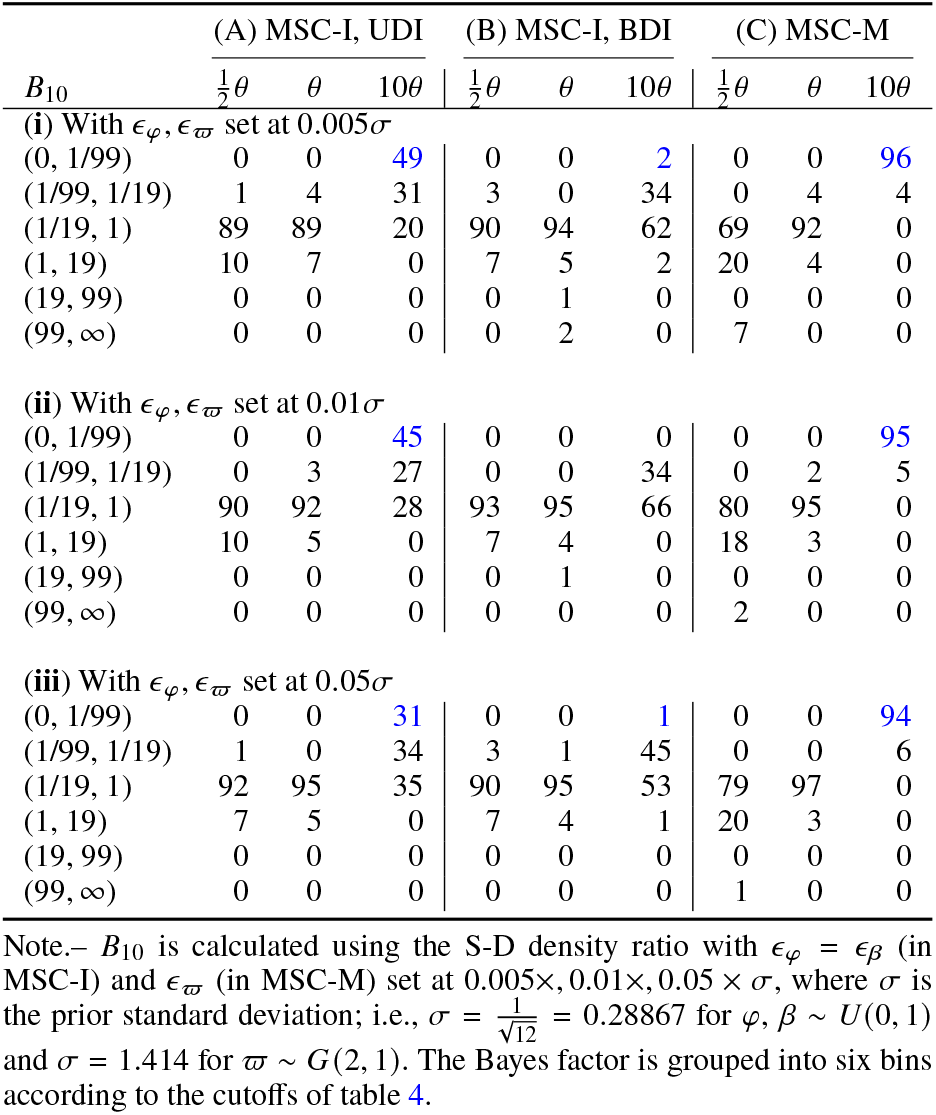
Bayes factor (*B*_10_) for testing introgression in 100 replicate datasets simulated under the null model of no gene flow (*H*_0_, MSC, Fig. 4A), and analyzed under models of gene flow (*H*_1_, MSC-I and MSC-M, Fig. 4B–D)

| $B_{10}$ | (A) MSC-I, UDI | | | (B) MSC-I, BDI | | | (C) MSC-M | | |
| --- | --- | --- | --- | --- | --- | --- | --- | --- | --- |
| | $\frac{1}{2}\theta$ | $\theta$ | $10\theta$ | $\frac{1}{2}\theta$ | $\theta$ | $10\theta$ | $\frac{1}{2}\theta$ | $\theta$ | $10\theta$ |
| (i) With $\epsilon_\varphi, \epsilon_\varpi$ set at $0.005\sigma$ | | | | | | | | | |
| (0, 1/99) | 0 | 0 | 49 | 0 | 0 | 2 | 0 | 0 | 96 |
| (1/99, 1/19) | 1 | 4 | 31 | 3 | 0 | 34 | 0 | 4 | 4 |
| (1/19, 1) | 89 | 89 | 20 | 90 | 94 | 62 | 69 | 92 | 0 |
| (1, 19) | 10 | 7 | 0 | 7 | 5 | 2 | 20 | 4 | 0 |
| (19, 99) | 0 | 0 | 0 | 0 | 1 | 0 | 0 | 0 | 0 |
| (99, ∞) | 0 | 0 | 0 | 0 | 2 | 0 | 7 | 0 | 0 |
| (ii) With $\epsilon_\varphi, \epsilon_\varpi$ set at $0.01\sigma$ | | | | | | | | | |
| (0, 1/99) | 0 | 0 | 45 | 0 | 0 | 0 | 0 | 0 | 95 |
| (1/99, 1/19) | 0 | 3 | 27 | 0 | 0 | 34 | 0 | 2 | 5 |
| (1/19, 1) | 90 | 92 | 28 | 93 | 95 | 66 | 80 | 95 | 0 |
| (1, 19) | 10 | 5 | 0 | 7 | 4 | 0 | 18 | 3 | 0 |
| (19, 99) | 0 | 0 | 0 | 0 | 1 | 0 | 0 | 0 | 0 |
| (99, ∞) | 0 | 0 | 0 | 0 | 0 | 0 | 2 | 0 | 0 |
| (iii) With $\epsilon_\varphi, \epsilon_\varpi$ set at $0.05\sigma$ | | | | | | | | | |
| (0, 1/99) | 0 | 0 | 31 | 0 | 0 | 1 | 0 | 0 | 94 |
| (1/99, 1/19) | 1 | 0 | 34 | 3 | 1 | 45 | 0 | 0 | 6 |
| (1/19, 1) | 92 | 95 | 35 | 90 | 95 | 53 | 79 | 97 | 0 |
| (1, 19) | 7 | 5 | 0 | 7 | 4 | 1 | 20 | 3 | 0 |
| (19, 99) | 0 | 0 | 0 | 0 | 0 | 0 | 0 | 0 | 0 |
| (99, ∞) | 0 | 0 | 0 | 0 | 0 | 0 | 1 | 0 | 0 |

Note.— $B_{10}$ is calculated using the S-D density ratio with $\epsilon_\varphi = \epsilon_\beta$ (in MSC-I) and $\epsilon_\varpi$ (in MSC-M) set at $0.005\times, 0.01\times, 0.05\times\sigma$ , where $\sigma$ is the prior standard deviation; i.e., $\sigma = \frac{1}{\sqrt{12}} = 0.28867$ for $\varphi, \beta \sim U(0, 1)$ and $\sigma = 1.414$ for $\varpi \sim G(2, 1)$ . The Bayes factor is grouped into six bins according to the cutoffs of table 4.

The Bayes factor *B*_10_ may be calibrated with reference to posterior model probabilities ℙ (*H*_1_ | ***x***) (table 4). At the 5% level (or *B*_10_ > 19), the false positive rate of the test is 0 for all but two settings, where the rate is 1% and 2%, lower than the nominal 5% (table 3). Here we are examining a frequentist property (the type-I error rate) of a Bayesian test, so one does not expect a match between the error rate and the significance level. Nevertheless, the false-positive rate of the Bayesian test is noted to be well below the nominal significance level.

**Table 4:**
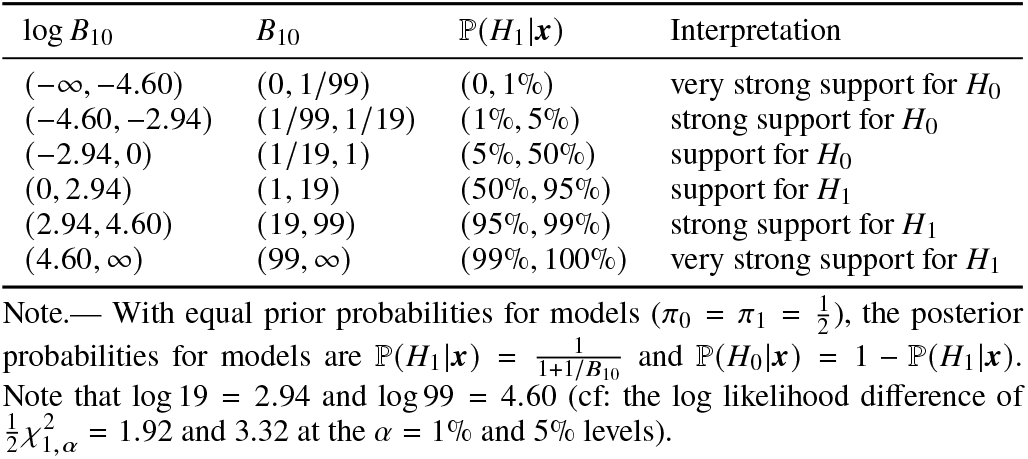
Calibration of the Bayes factor via posterior model probabilities.

**Table 5:**
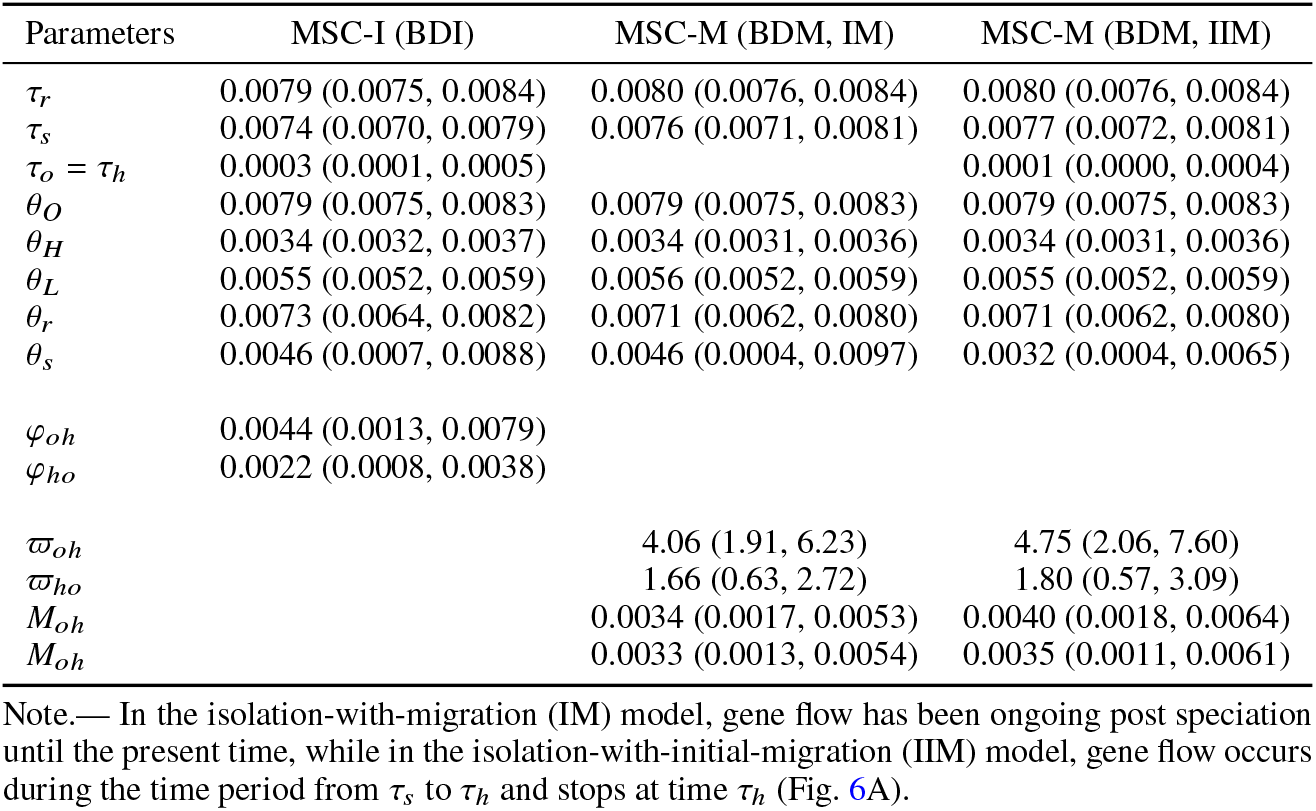
Posterior means and 95% HPD CIs for parameters under the BDI and BDM models obtained from bpp analyses of the lizard data.

| Parameters | MSC-I (BDI) | MSC-M (BDM, IM) | MSC-M (BDM, IIM) |
| --- | --- | --- | --- |
| $\tau_r$ | 0.0079 (0.0075, 0.0084) | 0.0080 (0.0076, 0.0084) | 0.0080 (0.0076, 0.0084) |
| $\tau_s$ | 0.0074 (0.0070, 0.0079) | 0.0076 (0.0071, 0.0081) | 0.0077 (0.0072, 0.0081) |
| $\tau_o = \tau_h$ | 0.0003 (0.0001, 0.0005) | | 0.0001 (0.0000, 0.0004) |
| $\theta_O$ | 0.0079 (0.0075, 0.0083) | 0.0079 (0.0075, 0.0083) | 0.0079 (0.0075, 0.0083) |
| $\theta_H$ | 0.0034 (0.0032, 0.0037) | 0.0034 (0.0031, 0.0036) | 0.0034 (0.0031, 0.0036) |
| $\theta_L$ | 0.0055 (0.0052, 0.0059) | 0.0056 (0.0052, 0.0059) | 0.0055 (0.0052, 0.0059) |
| $\theta_r$ | 0.0073 (0.0064, 0.0082) | 0.0071 (0.0062, 0.0080) | 0.0071 (0.0062, 0.0080) |
| $\theta_s$ | 0.0046 (0.0007, 0.0088) | 0.0046 (0.0004, 0.0097) | 0.0032 (0.0004, 0.0065) |
| $\varphi_{oh}$ | 0.0044 (0.0013, 0.0079) | | |
| $\varphi_{ho}$ | 0.0022 (0.0008, 0.0038) | | |
| $\varpi_{oh}$ | | 4.06 (1.91, 6.23) | 4.75 (2.06, 7.60) |
| $\varpi_{ho}$ | | 1.66 (0.63, 2.72) | 1.80 (0.57, 3.09) |
| $M_{oh}$ | | 0.0034 (0.0017, 0.0053) | 0.0040 (0.0018, 0.0064) |
| $M_{oh}$ | | 0.0033 (0.0013, 0.0054) | 0.0035 (0.0011, 0.0061) |

Note.— In the isolation-with-migration (IM) model, gene flow has been ongoing post speciation until the present time, while in the isolation-with-initial-migration (IIM) model, gene flow occurs during the time period from $\tau_s$ to $\tau_h$ and stops at time $\tau_h$ (Fig. 6A).

Note that the Bayesian test makes a ‘stronger’ inference than the LRT in that it can strongly reject *H*_1_ and support *H*_0_ whereas the LRT may fail to reject *H*_0_ but will never strongly support it. At the 5% level (*B*_10_ < 0.0526), *H*_1_ is rejected in 72% and 100% of datasets simulated at deep divergence (*τ* = 10*θ*) when the alternative hypothesis is MSC-I UDI and MSC-M, respectively (table 3). Strong support for *H*_0_ rarely occurs at shallow divergence 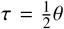, suggesting that the data are far more informative at deep divergence. These results are consistent with the simulation of Thawornwattana *et al*. (2026).

Posterior means and the 95% highest posterior density (HPD) credible intervals (CIs) for parameters in the three alternative models of gene flow (UDI, BDI, and MSC-M; Fig. 4B–D) are summarized in Figure S4. We first consider the UDI model. The population size parameter *θ* is well estimated, but the other parameters (*τ*_*R*_, *τ*_*Y*_ , *φ*_*Y*_) are not. In particular the introgression probability *φ*_*Y*_ has CIs that cover nearly the whole range (0, 1). While wide CIs might indicate that the data are uninformative about gene flow, here a stronger conclusion may be more appropriate. Note that the summaries in Figure S4 do not accommodate the fact that all of the three segments a, b, and c of Figure 1D represent the scenario of no gene flow. For example, both *φ* = 0 and *φ* = 1, as well as any value for *φ* (when *τ*_*Y*_ = *τ*_*R*_), represent the true model of no gene flow and are thus plausible, but the posterior mean and the HPD CI of *φ* generated from the MCMC sample are implausible. Similarly under the BDI model the neighborhoods of the five segments/regions that represent the null model of no gene flow (table 1(ii)) represent the same biological scenario of no gene flow, and this fact is not accommodated in the summaries for *φ*_*X*_ and *φ*_*Y*_ in Figure S4.

As discussed above, the model of gene flow is rejected in many of these datasets, especially at deep divergence (table 3). In such cases, the correct conclusion is strong evidence against gene flow (if *B*_10_ < 0.01 or 0.05), rather than a lack of information in the data. We suggest that if the estimated introgression probability between sister lineages has large CIs, one should rely on the Bayesian test to assess the evidence for gene flow, as discussed in the paper.

The MSC-M model (the IM model) does not involve the irregularities of the MSC-I models discussed in this paper. Posterior summaries of parameters under the model (*θ, τ*_*R*_, *M*) are all well-behaved and easy to interpret (Fig. S4). In particular, estimates of *M* are close to 0 (the true value), supporting the null model of no gene flow. The data are far more informative (with better estimates of *M*) on the deep tree than on the shallow trees.

### Bayesian test of gene flow between sister species has high power

Results of the Bayesian test when data were simulated under the MSC-I and MSC-M models of Figure 4B&C, with *A* → *B* (or *X* → *Y*) gene flow are summarized in Figure 7. First we consider data simulated under UDI (Fig. 7A–C). When the data are analyzed under the correct UDI model (I-UDI), the test has ∼100% power except for the very shallow tree 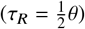 and low rates of gene flow (*φ* ≤ 0.1). For data simulated under UDI, the BDI model is over-parametrized but not misspecified. The test under the BDI showed similar performance to the use of the UDI model. We implemented the MSC-M model under two priors, *ϖ* ∼ *G* (2, 0.01) with mean 200 and *ϖ* ∼ *G* (2, 1) with mean 2. The test using the MSC-M model, in particular, under the *ϖ* ∼ *G* (2, 1) prior (Fig. 7C^′^, I-M) had lower power than under UDI or BDI. In particular, power dropped when the introgression probability is very high. We note that when 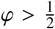, most sequences from *B* are traced to the introgression parent *A*. It should be noted that in all datasets and all models, *B*_10_ > 1 (Fig. 7A–C), suggesting that the Bayesian test supports gene flow, even if the support may not reach the cutoff of *B*_10_ > 19 or 99.

**Fig. 7:**
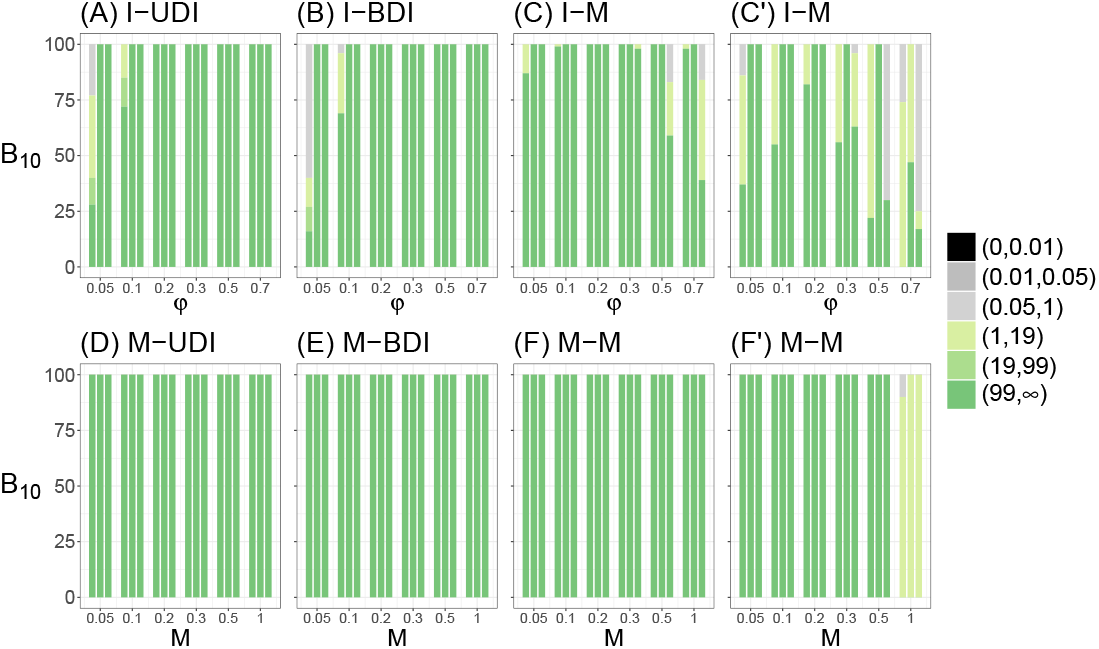
Power of the Bayesian test of gene flow between sister lineages. Bayes factor (*B*_10_) calculated using data simulated under the MSC-I or MSC-M models of figure 4**B**&**C**, and analyzed under the three models of figure 4**B**–**D**. The setting is in the format of simulation model-analysis model: for example, in M-UDI, data are simulated under MSC-M and analyzed under MSC-I (UDI) (Fig. 4**B**&**C**). The MSC-M model was implemented under two priors, *ϖ* ∼ *G* (2, 0.01) with mean 200 in panels **C & F**, and *ϖ* ∼ *G* (2, 1) with mean 2 in panels **C**^′ &^ **F**^′^. The stacked bar plots show the proportions of datasets (out of 100 replicates) in which *B*_10_ falls in different intervals: 0–0.01 (strong rejection of gene flow), 0.01–0.05 (rejection of gene flow), 0.05–1 (preference for no gene flow), 1–19 (preference for gene flow), 19–99 (support for gene flow), and 99–∞ (strong support for gene flow). The three bars in each group correspond to three divergence levels, with 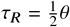 (very shallow), *θ* (shallow), and 10*θ* (deep). *B*_10_ is calculated via the S-D density ratio with *ϵ*_*φ*_, *ϵ*_*β*_ in MSC-I and *ϵ* in MSC-M set to 0.01*σ*, where *σ* is the prior standard deviation; use of 0.005*σ* and 0.05*σ* produces very similar results.

Next we consider data simulated under the MSC-M model (Fig. 7D–F). The test has full power at the 1% level (with *B*_10_ > 99) under the two MSC-I models (M-UDI, M-BDI) for all three species split times and for all levels of migration rates. Misspecification of the mode of gene flow (i.e., the true model assumes continuous gene flow while the analysis model is discrete) has not had any major impact. The test under MSC-M had full power except for the combination of *M* = 1 and the *ϖ* ∼ *G* (2, 1) prior. We note that just as *φ* ≈ 1 in MSC-I represents the scenario of isolation with no gene flow, *M* = ∞ represents the scenario of panmixia, with no gene flow.

Overall there is a greater amount of gene flow in data simulated under the MSC-M model. The expected amount of introgression under the MSC-M model, measured by the probability that any sequence from species *B* is traced to population *A* (irrespective of the migration time), when one traces the genealogical history of the sampled sequences backward in time, is

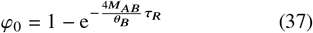

(Huang *et al*., 2022, eq. 10). At the low rate of *M* = 0.05, *φ*_0_ = 0.095, 0.181, 0.865 for the three divergence levels (Fig. S5), higher than the low rate *φ* = 0.05 in the MSC-I (UDI) model. This should explain why all tests have full power at the low rate in data simulated under MSC-M. At the high rate of *M* = 1, we have *φ*_0_ = 0.865, 0.982, 1.000, higher than *φ* = 0.7 in the UDI model. Data simulated under the MSC-M model with *φ*_0_ ≈ 1 may be expected to resemble data simulated under UDI with *φ* = 1 (which is equivalent to the MSC model of no gene flow; see Fig. 1C). This may explain the loss of power of the Bayesian test to detect gene flow at very high migration rates (*M*).

Compared with the true values used in the simulation, *ϖ* = 40, 80, 160, 240, 400 and 800 at the six values of *M*, the prior *ϖ* ∼ *G* (2, 1) has a mean that is far too small, creating conflicts between the prior and the data and leading to reduced power (Fig. 7D–F^′^). In practical data analysis, *G* (2, 0.1) with the prior mean 20 may be a good choice.

### Bayesian estimation of the rate of introgression between sister species is reliable only at deep species divergences

We examine the precision and accuracy of parameter estimation in data analyzed in Figure 7, simulated under models with gene flow. In particular, we are interested in the rate of gene flow (*φ* under MSC-I and *M* under MSC-M; Fig. 4B–D). The results are summarized in Figure 8.

**Fig. 8:**
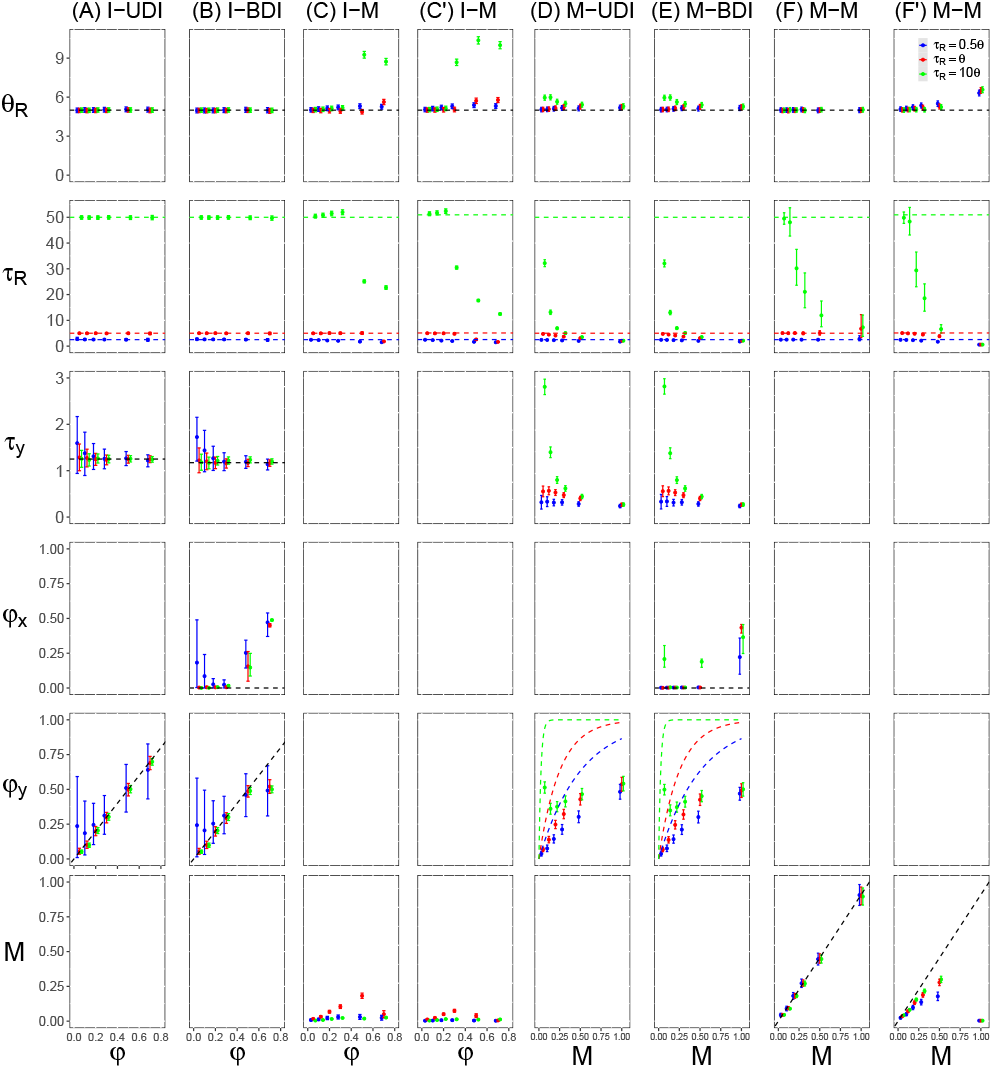
Average posterior means and 95% HPD CIs of parameters in bpp analysis of data of figure 7. See legend to figure 7. Dotted lines in the plots for *φ*_*y*_ in the M-UDI and M-BDI settings represent the expected amount of gene flow in the true MSC-M model (*φ*_0_ in eq. 37).

First we consider data simulated under the UDI model (Fig. 8A–C). Under the correct UDI model, species split time *τ*_*R*_ and the population size (*θ*) are well estimated. Note that we assumed the same population size for all species on the tree, so there is a huge amount of information in the data about *θ*. The introgression time *τ*_*Y*_ and the introgression probability *φ* involved large uncertainties when the true *φ* is small (< 0.2, say). When *φ* is very small, the null model of no gene flow is nearly correct, and the three segments of Figure 1C (where *φ* = 0, 1, or any value between 0–1) are expected to fit the data nearly equally well; in other words, there exist ridges in the posterior surface close to the three segments of Figure 1C, and the data are not informative about *φ*. The poor estimation of *τ*_*Y*_ and *φ* form a sharp contrast with the precise and accurate estimation of *τ*_*R*_ and *θ*. Furthermore, the species split time has a large impact, with the CIs for *φ* to be much narrower for the deep tree (with *τ*_*R*_ = 10*θ*) than for the very shallow tree (with 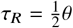) (Fig. 8).

The analysis under BDI (Fig. 8B, I-BDI) produces similar estimates of the species split time and introgression time (*τ*_*R*_, *τ*_*Y*_) to those under UDI, but the estimates of introgression probabilities (*φ*_*X*_, *φ*_*Y*_) show interesting patterns, different from the analysis under UDI (the I-UDI setting). For data generated at low *φ* in the true UDI model, both *φ*_*X*_ and *φ*_*Y*_ are poorly estimated. This is similar to the poor estimation of *φ*_*Y*_ under UDI, due to the issue with the summary: at the low rate, the null model of no gene flow is nearly correct, and the four lines and a plane under the BDI model (eq. 34, table 1(ii)), where *φ*_*X*_ and *φ*_*Y*_ are either 0 or 1 or anywhere over (0, 1), are expected to fit the data nearly equally well. At high *φ* in the true UDI model, the BDI model underestimates *φ*_*Y*_ (which has the true value *φ*) and overestimates *φ*_*X*_ (which has the true value 0), having difficulty in determining the direction of introgression. This may reflect the label-switching unidentifiability problem for the BDI model, as discussed by Yang and Flouri (2022). Under the BDI model, without constraints on *θ*, two sets of parameter values, ***θ*** = (*φ*_*X*_, *φ*_*Y*_ , *θ*_*X*_, *θ*_*Y*_) and ***θ***^′^ = (1 − *φ*_*X*_, 1 − *φ*_*Y*_ , *θ*_*Y*_ , *θ*_*X*_) , are unidentifiable (Yang and Flouri, 2022, Fig. 1). Similar unidentifiability involving *φ*_*X*_ and *φ*_*Y*_ occurs when all populations are assumed to have the same size (*θ*).

Next we consider data simulated under the MSC-M model (Fig. 8D–F). Estimates under UDI and BDI at shallow divergence 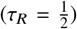 and low migration rate (*M*) are very poor. As discussed above, when the null model of no gene flow is nearly correct, our summaries in Figure 8 do not account for the fact that different parts of the parameter space represent the same biological scenario of no gene flow. When the migration rate is higher, the estimates improve, in particular at high divergence. However, when *M* is very high, the model will again be close to the null model of no gene flow. In general the MSC-I model does not recover all the gene flow that has occurred. The results are similar to previous simulations in which data were generated under the MSC-M model and analyzed under the MSC-I model (Huang *et al*., 2022, *φ* against *M* under the IM model).

In the M-M setting, the MSC-M model with *A* → *B* migration is used to both simulate and analyze the data (Fig. 8D–F). This case shows a complex pattern with respect to the species split time (*τ*_*R*_) and the migration rate *M*. When *M* is low (0.05 or 0.1), both *θ* and *τ* are well estimated, as is the migration rate *M*. When trees only 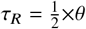, but poor for the deep tree. The 0.2 ≤ *M* ≤ 0.5, the estimates are good for the shallow prior on *ϖ* affected the posterior estimate (Fig. 8F&F^′^). Note that at *M* ≥ 0.2, *φ*_0_ = 1.000 on the deep tree (Fig. S5), and at such extreme amount of gene flow, the estimate of *τ*_*R*_ may collapse.

### Asymptotic analysis to assess the impact of species split time

To understand the effect of species split time on the information content in the data concerning the introgression probability *φ* under the UDI model (Fig. 4B), we consider the simple case of only two sequences per locus, with one sequence per species. The large-sample behavior of the estimation method (when the number of loci *L* → ∞) is analytically tractable (Huang *et al*., 2022). We assume one *θ* for all populations, so there are 4 parameters in the model, ***θ*** = (*φ, τ*_*Y*_ , *τ*_*R*_, *θ*). Let 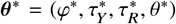 be the true parameter values. The alignment at each locus is summarized as the observed number (*x*) of differences at *n* sites. Let the probability of observing *x* differences at *n* sites be *p* (*x*; ***θ***), given by Huang *et al*. (2022, eq. S9). When the number of loci *L* → ∞, the per-locus log likelihood becomes

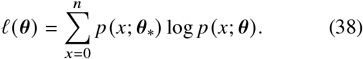

Note that here *n* is the sequence length so that the number of variable sites (*x*) has the range (0, *n*).

We fix the population size parameter *θ* at the true value, as it is typically well estimated. For given *τ*_*Y*_ and *φ*, we estimate *τ*_*R*_ by maximizing ℓ, and calculate the optimized ℓ as a function of the other two parameters (Fig. 9). We used three values for 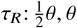, and 10*θ*, and three values for *φ*: 0.01, 0.1, and 0.6. The log-likelihood contour for *τ*_*Y*_ and *φ* is shown in Figure 9.

**Fig. 9:**
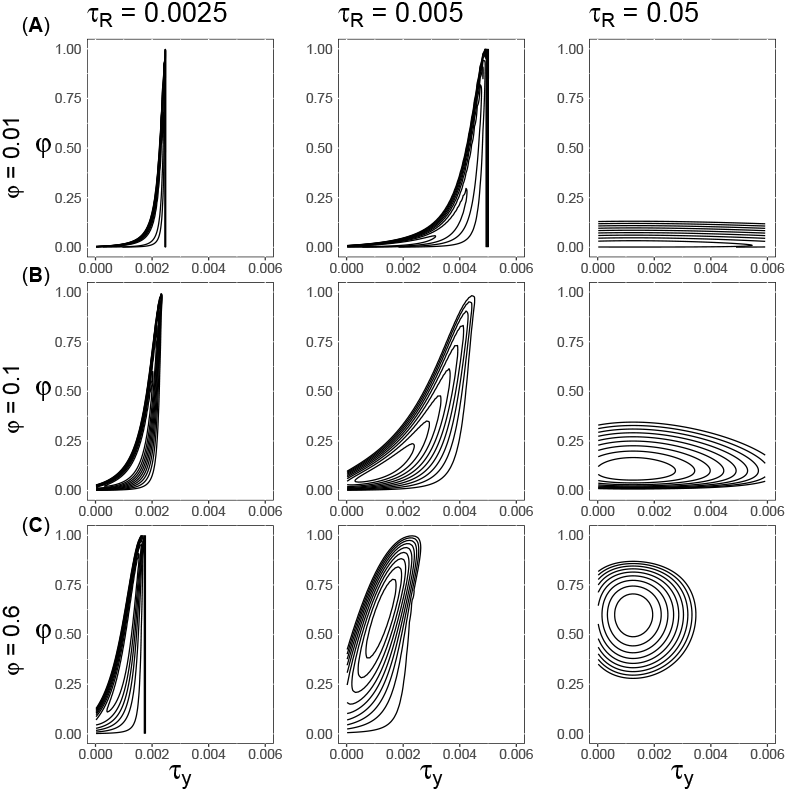
Contour plot of the log likelihood (eq. 38) as a function of *τ*_*Y*_ and *φ* under the MSC-I UDI model (Fig. 4**B**) for the limiting case of *L* = ∞ loci, generated using the true parameter values *φ* = 0.01, 0.1 or 0.6, and 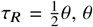, or 10*θ*, with *θ* = 0.005 and 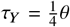. Each locus consists of two sequences (one from each species), summarized as the number of differences at *n* sites. Here *θ* = 0.005 is fixed at the true value, while *τ*_*R*_ is optimized given *φ* and *τ*_*Y*_ ; fixing *τ*_*R*_ at the true value produces nearly identical results.

When *φ* = 0.01 and the true *τ*_*R*_ = ^1^ *θ*, the log-likelihood surface has an elongated foot shape, similar to the Bayesian posterior surface for the two parameters of Figure 1F. Information content on *φ* improved when the species split time is larger (*τ*_*R*_ = 10*θ*) or when the rate of gene flow is higher (*φ* = 0.1 or 0.6). In particular, as *φ* increased, the contour lines became more circular, resulting in narrower CIs for both parameters (Fig. 9).

In the case of two sequences per locus and treating the coalescent time as data, Thawornwattana *et al*. (2023, eq. 2) give the Fisher information for estimating *φ*_*Y*_ as

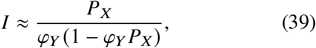

where 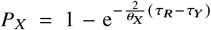 is the probability that two sequences (*a, b*) entering population *X* coalesce in population *X* rather than in population *R* (Fig. 4B). The data are more informative about *φ*_*Y*_ if *P*_*X*_ is larger (i.e, if species divergence is deeper with *τ*_*R*_ ≫ *τ*_*Y*_ and/or if population size *θ*_*X*_ is smaller).

### Inference of gene flow between Sceloporus lizard species

We analyzed a ddRAD dataset from three *Sceloporus* lizard species, *Sceloporus orcutti* (O), *S. hunsakeri* (H), and *S. licki* (L) to test for gene flow between the sister species *S. orcutti* and *S. hunsakeri* (Fig. 6A). We ran bpp under the bidirectional introgression (BDI) and migration (BDM) models. First we note that estimates of parameters shared between the two models were very similar (table 5). The introgression time under BDI model was nearly zero, indicating that gene flow between the two species may be ongoing or stopped very recently. The rate of gene flow was small, with *φ*_*oh*_ = 0.0044 (with the 95% HPD CI 0.0013–0.0079) and *φ*_*ho*_ = 0.0022 (0.0008–0.0038) (Fig. 6B, table 5). The BDM model assumes ongoing gene flow, with *M*_*oh*_ = *M*_*ho*_ = 0.0033 migrant individuals in each direction per generation (table 5). Those rates translate to *φ*_0,*o* →*h*_ = 1 − e^−4×0.003433/0.003388×0.007606^ = 0.0304, and *φ*_0,*h*→ *o*_ = 0.0125 according to eq. 37, suggesting that the migration model recovered more gene flow than the MSC-I model. We also fitted the IIM model, in which gene flow occurs post species split but stops at a later time. This produced nearly identical estimates to the IM model assuming ongoing gene flow (table 5). Overall the posterior estimates suggest a small but significant amount of gene flow in each direction, and gene flow is either ongoing or stopped a short time ago.

We then applied the Bayesian test. Under the BDI model, one can test *H*_0_ : *φ*_*oh*_ = 0, *φ*_*ho*_ = 0 against *H*_1_ : *φ*_*oh*_ ≠ 0, *φ*_*ho*_ ≠ 0 (table 1(ii)). Rejection of *H*_0_ means introgression in at least one direction. We used each of the five segments for which the condition on priors holds to calculate the Bayes factor, and all calculations led to strong support for gene flow (with *B* ≈ ∞).

Figure 6d illustrates the test of *H*_0_ : *φ*_*oh*_ = 0 against *H*_1_ : *φ*_*oh*_ ≠ 0, with *φ*_*ho*_ treated a nuisance parameter (present in both *H*_0_ and *H*_1_). There exist three segments, each of which can be used to calculate *B*_10_, as described in table 1(i). Here we used the first segment, with 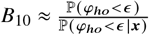. The data supported the *h* → *o* introgression, but the introgression probability was very low. However, the low introgression probabilities caused challenges to the S-D approach as *B* was sensitive to the choice of *ϵ*. The test under the BDM model (the IM model) does not involve irregularities. In Figure 6E, we used the same MCMC sample under BDM for Figure 6C to calculate the Bayes factor for testing three null hypotheses. The first is *H*_0_ : *ϖ*_*oh*_ = 0, against *H*_1_ : *ϖ*_*oh*_ > 0, with *ϖ*_*ho*_ treated as a nuisance parameter. The Bayes factor is given as 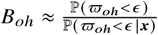, where ℙ (*ϖ*_*oh*_ < *ϵ* |***x***) was estimated by the proportion of MCMC samples in which *ϖ*_*oh*_ < *ϵ* while the prior probability *α* = ℙ (*ϖ*_*oh*_ < *ϵ*)= *G* (*ϵ* ; 2, 0.1) , the CDF of the gamma distribution. In fact we fix *α* = 0.001, 0.005 and obtain *ϵ* = 0.454, 1.035 via the quantile function. In the second test we have

*H*_0_ : *ϖ*_*ho*_ = 0 against *H*_1_ : *ϖ*_*ho*_ > 0. The third test is for *H*_0_ : *ϖ*_*oh*_ = 0, *ϖ*_*ho*_ = 0 against *H*_1_ : *ϖ*_*oh*_ > 0, *ϖ* > 0. We have 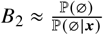, calculated by defining ∅ = (*ϖ*_*oh*_ < *ϵ* , *ϖ*_*ho*_ < *ϵ*). The three Bayes factors (*B*_*oh*_, *B*_*ho*_, *B*_2_) are plotted against the tail probability *α* in Figure 6E. All of them are ≫ 1, strongly supporting gene flow although there is a lack of precision when *B* is very large.

## Discussion

### Bayesian testing of gene flow between sister lineages and calculation of the Bayes factor

Considered a problem of statistical hypothesis testing (Jeffreys, 1935), the test of introgression between sister lineages involves a whole suite of challenging non-standard features, including boundary problems (the null value *φ* = 0 is at the boundary of the parameter space for *H*_1_), indeterminate parameters (e.g., when the introgression probability *φ* = 0, the time of introgression becomes unidentifiable), and in particular multiple routes from the alternative hypothesis to the null hypothesis (e.g., Fig. 1C). While these do not invalidate the definition and use of the Bayes factor for hypothesis testing, they pose challenges to the use of the S-D density ratio to calculate the Bayes factor. In this study, we show that each route from *H*_1_ to *H*_0_ can be used to calculate the Bayes factor via the S-D density ratio. When the condition on priors is satisfied over all routes, a combined estimate using all routes is also possible. Note that one draws the correct conclusion if the calculated Bayes factor falls into the right interval, and high precision may not be very important (table 4).

Our simulation demonstrates that our theories can be implemented in practice, and the resulting Bayesian test has low false positives and high power for testing gene flow between sister lineages.

For parameter estimation, we see that the posterior surface under introgression (MSC-I) models may involve nearly flat ridges when there is no gene flow or the rate of gene flow is very low. There are also label-switching unidentifiability issues under the bidirectional model (BDI) (Yang and Flouri, 2022). When the rate of introgression is low, one should exercise care when interpreting simple summaries of the MCMC samples (such as posterior means and CIs), and rely instead on the Bayes factor to assess the evidence for gene flow (Fig. S4).

Here we note a few advantages of the Bayes factor, relative to frequentist hypothesis testing such as the LRT (Yang, 2026). The Bayes factor penalizes parameter-rich models automatically. It works whether the compared models are nested or nonnested, and when the compared models have an irregular parameter space. The Bayes factor is also able to reject the alternative hypothesis with great force.

We note that many methods exist for calculating the marginal likelihood (see, for review, Fourment *et al*., 2020). Numerically stable algorithms, such as path-sampling or thermodynamic integration (Ogata, 1989; Gelman and Meng, 1998; Lartillot and Philippe, 2006), stepping-stones (Xie *et al*., 2011), and nested sampling (Skilling, 2006), require at least 50 runs of the MCMC algorithm for each of the two compared models (Fourment *et al*., 2020, table 1). The S-D approach to computing the Bayes factor requires only one run of the MCMC algorithm under the alternative model of gene flow (*H*_1_) (Dickey, 1971; Ji *et al*., 2023), and involves ∼ 100 times less computation than those sample-based methods, offering a major computational advantage. It is thus useful to extend the S-D approach to nonstandard conditions.

The irregularities discussed in this paper concern the test of gene flow between sister lineages under the discrete introgression models (UDI and BDI) and under the continuous IIM model (tables 1 & 2). Bayesian test of gene flow between non-sister lineages is not affected by those issues. Note also that the irregularities affect estimation of parameters for gene flow between sister species only when gene flow is absent or its rate is very low. At high rates or when there is clear evidence of gene flow and our objective is to estimate parameters such as the introgression probability, the irregularities are not relevant. When the MCMC samples the rest of the parameter space in *H*_1_, away from the three lines in Figure 1D, different points in the space represent distinct biological scenarios and are identifiable.

The theory developed in this paper applies as well if introgression is between sister lineages that are ancestral branches on the species phylogeny (Fig. 10). Whether one or both of the sister lineages involved are ancestral, there are three routes for reducing the UDI introgression model (*H*_1_) to the null model of no gene flow (*H*_0_). For example, in comparison of the two models of Figure 10A&B, the null point also consists of three lines or line segments, given by eq. 31 (Fig. 1C). The same holds true for the test comparing the two models of Figure 10C&D, in which case both sister lineages are ancestral. Similarly in the test under the BDI introgression model, there are five routes for reducing *H*_1_ to *H*_0_, as in the case where both involved species are extant (eq. 34).

**Fig. 10:**
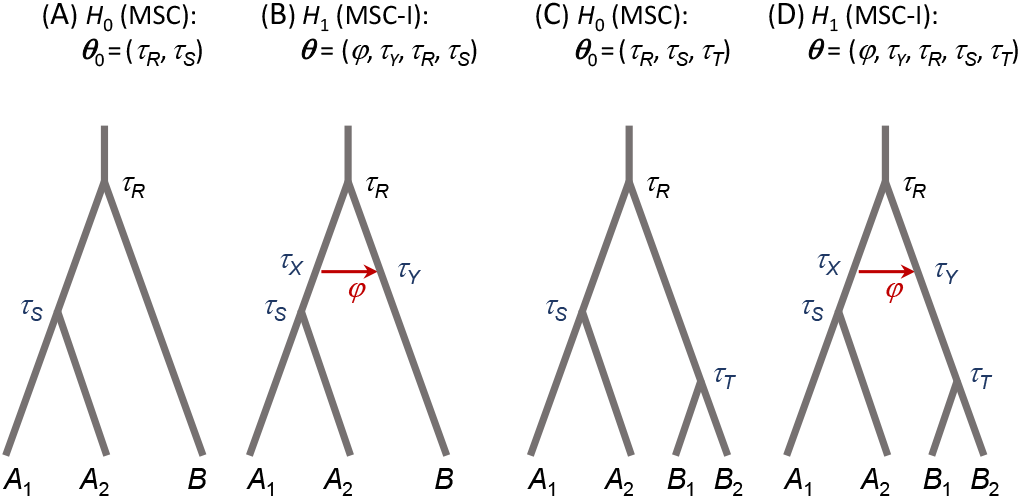
(**A, C**) Null (*H*_0_) and (**B, D**) alternative (*H*_1_) hypotheses for testing (unidirectional) introgression that involves one or two ancestral (sister) species. Population-size parameters (*θ*) are not shown.

### Alternative tests of gene flow between sister lineages

The irregularities of the testing problem discussed in this paper also impact frequentist hypothesis testing such as the LRT, and indeed often cause insurmountable difficulties for which no known solutions exist (Chernoff, 1954; Self and Liang, 1987; Mitchell *et al*., 2019; Brazzale and Mameli, 2024). In the case where *H*_0_ corresponds to *H*_1_ with one parameter of interest at the boundary of the space, the null distribution for the LRT statistic is a 50:50 mixture of 0 and χ^2^ (Chernoff, 1954). The problem of indeterminate parameters is harder, and in general the null distribution is unknown. The problem of multiple routes for reducing *H*_1_ to *H*_0_ does not appear to have been studied in the statistics literature.

In population genetics, the programs *∂*a*∂*i (Gutenkunst *et al*., 2009) and fastsimcoal2 (Excoffier *et al*., 2013) calculate a composite likelihood for data of joint site frequency spectrum (SFS). The likelihood is composite as the method ignores the lack of independence among linked SNP sites, or the lack of independence among population pairs when multiple populations are analyzed. The use of composite likelihood ignoring correlation and the use of data summaries that are not sufficient statistics make it hard to define the null distribution of the test statistic; even without the irregularities discussed here, the usual χ^2^ distribution may not apply. In practice, nonparametric resampling methods such as the bootstrap are used instead (Excoffier *et al*., 2013).

## Supplementary Material and Data Availability

Supplemental information is available at https://doi.org/10.5281/zenodo.xxxxxxxx. Supplementary data (sequence alignments and control files for running bpp are available through the Dryad Digital Repository at 10.5061/dryad.zkh1893pp.

## Acknowledgments

We thank Dr Adam Leaché for providing the spiny lizard data, and Drs Mengyu Li and Jun Liu for bringing to our attention Rousseau and Mengersen (2011). We are indebted to three anonymous reviewers for many constructive comments and suggestions.

## References

Blischak, P. D., Chifman, J., Wolfe, A. D., and Kubatko, L. S. 2018. HyDe: a Python package for genome-scale hybridization detection. Syst. Biol., 67(5): 821–829.

Brazzale, A. R. and Mameli, V. 2024. Likelihood asymptotics in nonregular settings: A review with emphasis on the likelihood ratio. Statist. Sci., 39: 322–345.

Chernoff, H. 1954. On the distribution of the likelihood ratio. Ann. Math. Stat., 25: 573–578.

Costa, R. J. and Wilkinson-Herbots, H. 2017. Inference of gene flow in the process of speciation: An efficient maximum-likelihood method for the isolation-with-initial-migration model. Genetics, 205(4): 1597–1618.

Dalquen, D., Zhu, T., and Yang, Z. 2017. Maximum likelihood implementation of an isolation-with-migration model for three species. Syst. Biol., 66: 379–398.

Dickey, J. M. 1971. The weighted likelihood ratio, linear hypotheses on normal location parameters. Ann. Math. Statist., 42(1): 204–223.

Excoffier, L., Dupanloup, I., Huerta-Sanchez, E., Sousa, V. C., and Foll, M. 2013. Robust demographic inference from genomic and SNP data. PLoS Genet., 9(10):e1003905.

Flouri, T., Jiao, X., Rannala, B., and Yang, Z. 2020. A Bayesian implementation of the multispecies coalescent model with introgression for phylogenomic analysis. Mol. Biol. Evol., 37(4): 1211–1223.

Flouri, T., Huang, J., Jiao, X., Kapli, P., Rannala, B., and Yang, Z. 2022. Bayesian phylogenetic inference using relaxed-clocks and the multispecies coalescent. Mol. Biol. Evol., 39(8):msac161.

Flouri, T., Jiao, X., Huang, J., Rannala, B., and Yang, Z. 2023. Efficient Bayesian inference under the multispecies coalescent with migration. Proc. Nat. Acad. Sci. U.S.A., 120(44):e2310708120.

Fourment, M., Magee, A. F., Whidden, C., Bilge, A., Matsen, F. A., and Minin, V. N. 2020. 19 dubious ways to compute the marginal likelihood of a phylogenetic tree topology. Syst. Biol., 69(2): 209–220.

Gelman, A. and Meng, X. 1998. Simulating normalizing constants: From importance sampling to bridge sampling to path sampling. Stat. Sci., 13: 163–185.

Gottscho, A. D., Hollingsworth, B. D., Espinal, J. L., Leaché, A. D., Reeder, T. W., and de Queiroz, K. 2025. Comparative phylogeography of lizards (squamata: Phrynosomatidae) in Baja California and expansion of Callisaurus draconoides within the North American deserts. J. Biogeography, page 10.1101/2025.04.29.650089.

Green, R. E., Krause, J., Briggs, A. W., Maricic, T., Stenzel, U., Kircher, M., Patterson, N., Li, H., Zhai, W., Fritz, M. H., Hansen, N. F. and Durand, E. Y., Malaspinas, A. S., Jensen, J. D., Marques-Bonet, T., Alkan, C., Prufer, K., Meyer, M., Burbano, H. A., Good, J. M., Schultz, R., Aximu-Petri, A., Butthof, A., Hober, B., Hoffner, B., Siegemund, M., Weihmann, A., Nusbaum, C., Lander, E. S., Russ, C., Novod, N., Affourtit, J., Egholm, M., Verna, C., Rudan, P., Brajkovic, D., Kucan, Z., Gusic, I., Doronichev, V. B., Golovanova, L. V., Lalueza-Fox, C., de la Rasilla, M., Fortea, J., Rosas, A., Schmitz, R. W., Johnson, P. L., Eichler, E. E., Falush, D., Birney, E., Mullikin, J. C., Slatkin, M., Nielsen, R., Kelso, J., Lachmann, M., Reich, D., and Paabo, S. 2010. A draft sequence of the Neandertal genome. Science, 328: 710–722.

Gutenkunst, R. N., Hernandez, R. D., Williamson, S. H., and Bustamante, C. D. 2009. Inferring the joint demographic history of multiple populations from multidimensional SNP frequency data. PLoS Genet, 5(10):e1000695.

Hey, J., Chung, Y., Sethuraman, A., Lachance, J., Tishkoff, S., Sousa, V. C., and Wang, Y. 2018. Phylogeny estimation by integration over isolation with migration models. Mol. Biol. Evol., 35(11):2805—-2818.

Hibbins, M. S. and Hahn, M. W. 2022. Phylogenomic approaches to detecting and characterizing introgression. Genetics, 220(2):iyab173.

Huang, J., Flouri, T., and Yang, Z. 2020. A simulation study to examine the information content in phylogenomic datasets under the multispecies coalescent model. Mol. Biol. Evol., 37(11): 3211–3224.

Huang, J., Thawornwattana, Y., Flouri, T., Mallet, J., and Yang, Z. 2022. Inference of gene flow between species under misspecified models. Mol. Biol. Evol., 39(12).

Jeffreys, H. 1935. Some tests of significance, treated by the theory of probability. Proc. Cam. Phil. Soc., 31: 203–222.

Ji, J., Jackson, D. J., Leaché, A. D., and Yang, Z. 2023. Power of Bayesian and heuristic tests to detect cross-species introgression with reference to gene flow in the Tamias quadrivittatus group of North American chipmunks. Syst. Biol., 72(2): 446–465.

Jiao, X., Flouri, T., and Yang, Z. 2021. Multispecies coalescent and its applications to infer species phylogenies and cross-species gene flow. Nat. Sci. Rev.

Jukes, T. H. and Cantor, C. R. 1969. Evolution of protein molecules. In Mammalian Protein Metabolism, pages 21–123. Academic Press, New York.

Kong, S., Swofford, D. L., and Kubatko, L. S. 2025. Inference of phylogenetic networks from sequence data using composite likelihood. Syst. Biol., 74(1): 53–69.

Lartillot, N. and Philippe, H. 2006. Computing Bayes factors using thermodynamic integration. Syst. Biol., 55: 195–207.

Lewis, P., Holder, M., and Holsinger, K. 2005. Polytomies and Bayesian phylogenetic inference. Syst. Biol., 54: 241–253.

Meng, C. and Kubatko, L. 2009. Detecting hybrid speciation in the presence of incomplete lineage sorting using gene tree incongruence: a model. Theo. Popul. Biol., 75(1): 35–45.

Mitchell, J. D., Allman, E. S., and Rhodes, J. A. 2019. Hypothesis testing near singularities and boundaries. Electron. J. Stat., 13(1): 2150–2193.

Ogata, Y. 1989. A Monte Carlo method for high dimensional integration. Numer. Math., 55: 137–157.

O’Hagan, A. and Forster, J. 2004. Kendall’s Advanced Theory of Statistics: Bayesian Inference. Arnold, London.

Payseur, B. A. and Rieseberg, L. H. 2016. A genomic perspective on hybridization and speciation. Mol. Ecol., 25(11): 2337–2360.

Rousseau, J. and Mengersen, K. 2011. Asymptotic behaviour of the posterior distribution in overfitted mixture models. J. R. Stat. Soc. B, 73: 689–710.

Roux, C., Fraisse, C., Romiguier, J., Anciaux, Y., Galtier, N., and Bierne, N. 2016. Shedding light on the grey zone of speciation along a continuum of genomic divergence. PLoS Biol., 14(12):e2000234.

Self, S. and Liang, K.-Y. 1987. Asymptotic properties of maximum likelihood estimators and likelihood ratio tests under nonstandard conditions. J. Am. Stat. Assoc., 82: 605–610.

Skilling, J. 2006. Nested sampling for general Bayesian computation. Bayesian Anal, 1: 833–859.

Solis-Lemus, C. and Ane, C. 2016. Inferring phylogenetic networks with maximum pseudolikelihood under incomplete lineage sorting. PLoS Genet., 12(3):e1005896.

Solis-Lemus, C., Bastide, P., and Ane, C. 2017. PhyloNetworks: a package for phylogenetic networks. Mol. Biol. Evol., 34(12): 3292–3298.

Sousa, V. and Hey, J. 2013. Understanding the origin of species with genome-scale data: modelling gene flow. Nat. Reviews Genet., 14(6): 404–414.

Suchard, M. A., Weiss, R. E., and Sinsheimer, J. S. 2003. Testing a molecular clock without an outgroup: derivations of induced priors on branch-length restrictions in a bayesian framework. Syst. Biol., 52: 48–54.

Thawornwattana, Y., Huang, J., Flouris, T., Mallet, J., and Yang, Z. 2023. Inferring the direction of introgression using genomic sequence data. Mol. Biol. Evol., 40(8):msad178.

Thawornwattana, Y., Flouris, T., Mallet, J., and Yang, Z. 2025. Inference of gene flow between species from genomic data when the mode, direction and lineages are misspecified. Mol. Biol. Evol., 42: 1–18.

Thawornwattana, Y., Rannala, B., and Yang, Z. 2026. On the robustness of Bayesian inference of gene flow to intragenic recombination and natural selection. Mol. Biol. Evol., page 10.1093/molbev/msaf327.

Verdinelli, I. and Wasserman, L. 1995. Computing bayes factors using a generalization of the savage-dickey density ratio. J. Am. Stat. Assoc., 90(430): 614–618.

Wen, D. and Nakhleh, L. 2018. Coestimating reticulate phylogenies and gene trees from multilocus sequence data. Syst. Biol., 67(3): 439–457.

Xie, W., Lewis, P. O., Fan, Y., Kuo, L., and Chen, M.-H. 2011. Improving marginal likelihood estimation for Bayesian phylogenetic model selection. Syst. Biol., 60: 150–160.

Yang, Z. 2007. Fair-balance paradox, star-tree paradox and Bayesian phylogenetics. Mol. Biol. Evol., 24: 1639–1655.

Yang, Z. 2026. Bayesian phylogenetic methods. In L. Knowels, editor, Encyclopaedia of Evolutionary Biology, pages 62–76. Elsevier, New York, 2nd edition.

Yang, Z. and Flouri, T. 2022. Estimation of cross-species introgression rates using genomic data despite model unidentifiability. Mol. Biol. Evol., 39(5):10.1093/molbev/msac083.

Yang, Z. and Rannala, B. 2005. Branch-length prior influences Bayesian posterior probability of phylogeny. Syst. Biol., 54: 455–470.

Yang, Z. and Rannala, B. 2010. Bayesian species delimitation using multilocus sequence data. Proc. Natl. Acad. Sci. U.S.A., 107: 9264–9269.

Yang, Z. and Zhu, T. 2018. Bayesian selection of misspecified models is overconfident and may cause spurious posterior probabilities for phylogenetic trees. Proc. Natl. Acad. Sci. U.S.A., 115(8): 1854–1859.

Yu, Y., Degnan, J. H., and Nakhleh, L. 2012. The probability of a gene tree topology within a phylogenetic network with applications to hybridization detection. PLoS Genet., 8(4):e1002660.

Zhu, T. and Yang, Z. 2012. Maximum likelihood implementation of an isolation-with-migration model with three species for testing speciation with gene flow. Mol. Biol. Evol., 29: 3131–3142.

