## Supplemental Information for "Bayesian test of gene flow between sister lineages using genomic data"

### SI text 1. Bayes factor for the two-sample test

Consider the two-sample test, in which samples are taken from two populations to test the equality of the population means with known variance. The data are $x_{ij} \sim N(\mu_i, 1)$, for $i = 1, 2$; $j = 1, \cdots, n$. The null hypothesis is $H_0 : \mu_1 = \mu_2 = \mu$ while the alternative hypothesis is $H_1 : \mu_1 \neq \mu_2$. The constraint $\Omega_0 : \mu_1 = \mu_2$ reduces $H_1$ to $H_0$. This test does not have any of the irregularities discussed in the paper. It is analytically tractable and serves to illustrate the S-D approach to calculating the Bayes factor and the use of reparametrization to deal with a null hypothesis formulated via equality constraints.

The data can be summarized as the two sample means, $\boldsymbol{x} = (\bar{x}_1, \bar{x}_2)$, with $\bar{x}_i = \frac{1}{n}\sum_j x_{ij}$, $i = 1, 2$. As $\bar{x}_1 - \bar{x}_2 \sim N(0, 2/n)$ under $H_0$, the likelihood ratio test statistic is

$$2\Delta\ell = \frac{n}{2}(\bar{x}_1 - \bar{x}_2)^2, \tag{S1}$$

to be compared with $\chi^2_1$.

For a Bayesian analysis, we assign the prior $\mu \sim N(0, 1/(2\tau))$ under $H_0$, and the prior

$$\begin{aligned} p(\mu_1, \mu_2) &= \phi(\mu_1; 0, 1/\tau)\phi(\mu_2; 0, 1/\tau) \\ &= \frac{\tau}{2\pi}\exp\Big\{-\frac{\tau}{2}(\mu_1^2 + \mu_2^2)\Big\} \end{aligned} \tag{S2}$$

under $H_1$, with $\tau = 1$.

The posterior of parameters under $H_1$ is

$$p(\mu_1, \mu_2 \mid \boldsymbol{x}) = \phi\Big(\mu_1; \tfrac{n\bar{x}_1}{\tau+n}, \tfrac{1}{\tau+n}\Big)\phi\Big(\mu_2; \tfrac{n\bar{x}_2}{\tau+n}, \tfrac{1}{\tau+n}\Big). \tag{S3}$$

The marginal likelihood is

$$\begin{aligned} m_0 &= p(\bar{x}_1, \bar{x}_2 \mid \Omega_0) \\ &= \int \phi(\mu; 0, \tfrac{1}{2\tau})\phi(\bar{x}_1; \mu, \tfrac{1}{n})\phi(\bar{x}_2; \mu, \tfrac{1}{n})\,\mathrm{d}\mu \\ &= \frac{n}{2\pi}\sqrt{\frac{\tau}{\tau+n}}\exp\Big\{\frac{n^2(\bar{x}_1 + \bar{x}_2)^2}{4(\tau+n)} - \frac{n(\bar{x}_1^2 + \bar{x}_2^2)}{2}\Big\} \\ &= \frac{n}{2\pi}\sqrt{\frac{\tau}{\tau+n}}\exp\Big\{\frac{-n^2(\bar{x}_1 - \bar{x}_2)^2 - 2\tau n(\bar{x}_1^2 + \bar{x}_2^2)}{4(\tau+n)}\Big\} \end{aligned} \tag{S4}$$

under $H_0$, and

$$\begin{aligned} m &= p(\bar{x}_1, \bar{x}_2) \\ &= \phi\Big(\bar{x}_1; 0, \tfrac{1}{\tau} + \tfrac{1}{n}\Big)\cdot\phi\Big(\bar{x}_2; 0, \tfrac{1}{\tau} + \tfrac{1}{n}\Big) \\ &= \frac{\tau n}{2\pi(\tau+n)}\exp\Big\{-\frac{\tau n}{2(\tau+n)}(\bar{x}_1^2 + \bar{x}_2^2)\Big\}. \end{aligned} \tag{S5}$$

under $H_1$. Thus the Bayes factor is

$$B_{10} = \frac{m}{m_0} = \sqrt{\frac{\tau}{\tau+n}}\exp\Big\{\frac{n^2(\bar{x}_1 - \bar{x}_2)^2}{4(\tau+n)}\Big\}. \tag{S6}$$

Note that under $H_0$, $\bar{x}_1 - \bar{x}_2 \sim N(0, \frac{2}{n})$, so that for large $n$, the Bayes factor has the form $B_{10} \approx c \cdot n^{-\frac{1}{2}}\,\mathrm{e}^{\frac{1}{2}\chi^2}$, where $c = \sqrt{\tau}$ is a constant and $\chi^2$ is a $\chi^2_1$ variable. When $H_0$ is true, $B_{10} \to 0$ at the rate $n^{-\frac{1}{2}}$.

To apply the S-D density ratio, we re-parametrize $H_1$ so that $\boldsymbol{\theta}' = (\mu_1, \delta)$, with $\delta = \mu_2 - \mu_1$. Then $\delta$ is the parameter of interest, with $H_0$ represented by $\delta = \delta_0 = 0$, while $\mu_1$ is the nuisance parameter. The prior under $H_1$ is given by eq. S2 through a change of variables as

$$p(\mu_1, \delta) = \frac{1}{2\pi/\tau}\exp\Big\{-\frac{\tau}{2}\big(\mu_1^2 + (\mu_1 + \delta)^2\big)\Big\}. \tag{S7}$$

The marginal prior for $\delta$ is

$$p(\delta) = \int p(\mu_1, \delta)\,\mathrm{d}\mu_1 = \frac{1}{\sqrt{4\pi/\tau}}\exp\Big\{-\frac{\tau}{4}\delta^2\Big\} \tag{S8}$$

or $\delta \sim N(0, 2/\tau)$.

To verify the condition on priors (eq. 1), note that from eq. S7 we have

$$p(\mu_1 \mid \delta) \propto \mathrm{e}^{-\frac{\tau}{2}(2\mu_1^2 + 2\mu_1\delta)}, \tag{S9}$$

or $p(\mu_1|\delta) = \phi(\mu_1; -\delta/2, 1/(2\tau))$. Thus $p(\mu_1|\delta_0) = \phi(\mu_1; 0, 1/(2\tau))$ under $H_1$ matches $p_0(\mu) = \phi(\mu; 0, 1/(2\tau))$ under $H_0$, and the condition holds.

From eq. S3, $\mu_1, \mu_2$ are independent in the posterior. Thus $\delta$ has the posterior

$$p(\delta \mid \boldsymbol{x}) = \phi\Big(\delta; \tfrac{n(\bar{x}_2 - \bar{x}_1)}{\tau+n}, \tfrac{2}{\tau+n}\Big). \tag{S10}$$

Using the prior and posterior of eqs. S8&S10, we get the Bayes factor via the S-D density ratio as

$$B_{10} = \frac{p(\delta_0)}{p(\delta_0 \mid x)} = \frac{\phi(0; 0, 2/\tau)}{\phi\Big(0; \tfrac{n(\bar{x}_2 - \bar{x}_1)}{\tau+n}, \tfrac{2}{\tau+n}\Big)}, \tag{S11}$$

which simplifies to eq. S6.

Let $(\mu_{1i}, \mu_{2i})$, $i = 1, \cdots, N$ be a sample from the posterior under $H_1$ (e.g., generated from an MCMC algorithm). Define $\Omega_\epsilon : |\mu_1 - \mu_2| < \epsilon$ to be the *null region*, a strip along the diagonal line $\mu_1 = \mu_2$. Then

$$B_{10} \approx \frac{\mathbb{P}(\Omega_\epsilon)}{\mathbb{P}(\Omega_\epsilon \mid \boldsymbol{x})} = \frac{\mathbb{P}\{|\mu_1 - \mu_2| < \epsilon\}}{\mathbb{P}\{|\mu_1 - \mu_2| < \epsilon \mid \boldsymbol{x}\}}, \tag{S12}$$

where $\mathbb{P}(\Omega_\epsilon|\boldsymbol{x})$ is the proportion of samples in which $|\mu_{1i} - \mu_{2i}| < \epsilon$. This should be $\approx p(\delta_0|\boldsymbol{x}) \cdot 2\epsilon$, with $p(\delta_0|\boldsymbol{x})$ from eq. S10. Similarly $\mathbb{P}(\Omega_\epsilon) = \mathbb{P}\big(|\mu_1 - \mu_2| < \epsilon\big) \approx p(\delta_0)2\epsilon$, with $p(\delta_0)$ from eq. S8. Thus $\mathbb{P}(\Omega_\epsilon)/\mathbb{P}(\Omega_\epsilon|\boldsymbol{x}) \approx p(\delta_0)/p(\delta_0|x)$, which is eq. S11.

### SI Text 2. The S-D density ratio for the one-sample test and the Borel-Kolmogorov paradox

Consider the one-sample test with unknown variance, in which a sample is taken to test $H_0 : N(0, \sigma^2)$ against $H_1 : N(\mu, \sigma^2)$, with $\sigma^2$ unknown. The parameter vector is $\boldsymbol{\theta}_0 = (\sigma^2)$ under $H_0$ and $\boldsymbol{\theta} = (\mu, \sigma^2)$ under $H_1$. The data are an i.i.d. sample of size $n$, and may be summarized as the sample mean $\bar{x}$ and variance $s^2 = \frac{1}{n}\sum(x_i - \bar{x})^2$, so that $\boldsymbol{x} = (\bar{x}, s^2)$. The likelihood functions under the two models are

$$L_0(\boldsymbol{\theta}_0) = (2\pi\sigma^2)^{-n/2} \exp\left\{-\frac{1}{2\sigma^2}[n\bar{x}^2 + ns^2]\right\},$$
$$L(\boldsymbol{\theta}) = (2\pi\sigma^2)^{-n/2} \exp\left\{-\frac{1}{2\sigma^2}[n(\bar{x}-\mu)^2 + ns^2]\right\}. \quad \text{(S13)}$$

Under $H_0$, we specify an inverse gamma prior, $\sigma^2 \sim \mathrm{IG}(\alpha, \beta)$. Under $H_1$, we let $\sigma^2 \sim \mathrm{IG}(\alpha, \beta)$, and then given $\sigma^2$, a Gaussian prior on $\mu$, with $\mu|\sigma^2 \sim N(0, k\sigma^2)$ with $k$ given, so that

$$p(\boldsymbol{\theta}) = \frac{\beta^\alpha}{\Gamma(\alpha)}(\sigma^2)^{-\alpha-1} \mathrm{e}^{-\beta/\sigma^2} \cdot \frac{1}{\sqrt{2\pi k\sigma^2}} \mathrm{e}^{-\frac{1}{2k\sigma^2}\mu^2}. \quad \text{(S14)}$$

The marginal likelihood values under $H_0$ and $H_1$ are

$$m_0 = (2\beta)^\alpha \frac{\Gamma(\alpha + \frac{n}{2})}{\Gamma(\alpha)} \pi^{-\frac{n}{2}} \left[2\beta + ns^2 + n\bar{x}^2\right]^{-\alpha-\frac{n}{2}},$$
$$m = (2\beta)^\alpha \frac{\Gamma(\alpha + \frac{n}{2})}{\Gamma(\alpha)} (1+nk)^{-\frac{1}{2}} \pi^{-\frac{n}{2}} \left[2\beta + ns^2 + \frac{n\bar{x}^2}{1+nk}\right]^{-\alpha-\frac{n}{2}} \quad \text{(S15)}$$

(O'Hagan and Forster, 2004, p.170).

The Bayes factor is

$$B = \frac{m}{m_0} = \frac{1}{\sqrt{1+nk}} \left[\frac{2\beta + ns^2 + n\bar{x}^2}{2\beta + ns^2 + \frac{n\bar{x}^2}{1+nk}}\right]^{\alpha+\frac{n}{2}} \quad \text{(S16)}$$

(O'Hagan and Forster, 2004, eq. 7.10; note that $\frac{a_1}{a_2}$ in the equation should be $\frac{a_2}{a_1}$)[1].

Note that under $H_0$, $\frac{\bar{x}}{\sqrt{s^2/(n-1)}} \sim t_{n-1}$, the $t$ distribution with $n-1$ degrees of freedom. Also $\left(1+\frac{a}{n}\right)^n \to \mathrm{e}^a$ when $n \to \infty$. Thus for large $n$, $t_{n-1}$ is approximately a $N(0,1)$ variable, and the Bayes factor has the form $B_{10} \approx c \cdot n^{-\frac{1}{2}} \mathrm{e}^{\frac{1}{2}\chi^2}$, where $c = k^{-\frac{1}{2}}$ is a constant and $\chi^2$ is a $\chi^2_1$ variable.

To apply the S-D approach, note that the conditional prior under $H_1$ is

$$p(\sigma^2|\mu = 0) \propto (\sigma^2)^{-\alpha-1-\frac{1}{2}} \mathrm{e}^{-\beta/\sigma^2}, \quad \text{(S17)}$$

which is the density for $\mathrm{IG}(\alpha+\frac{1}{2}, \beta)$. This does not match the prior $\sigma^2 \sim \mathrm{IG}(\alpha, \beta)$ under $H_0$.

[1] In Example 7.3 of O'Hagan and Forster (2004, p.169-171), $\mu$ is assigned the prior $N(m, w\sigma^2)$ under $H_1$ but here we use $\mu \sim N(0, k\sigma^2)$, with the prior mean fixed at 0. Symbols in that Example correspond to those used here as follows: $w \to k$, $b \to 2\alpha$, $a \to 2\beta$, and $s^2 \to ns^2$, and the marginal likelihood values are $f_1(x) \to m$, and $f_2(x) \to m_0$, with $\frac{f_1(x)}{f_2(x)}$ in eq. 7.11 to be the Bayes factor $B = \frac{m}{m_0}$ here. We note a few typos in Example 7.3: in eq. 7.9, $(\mu - \bar{x})^2$ should be $n(\mu - \bar{x})^2$; in the equation for $f_1(x)$ on p.170, $\Gamma(b)$ should be $\Gamma(\frac{1}{2}b)$; and in eq. 7.10, $(\frac{a_1}{a_2})$ should be $(\frac{a_2}{a_1})$.

Now consider an alternative parametrization of $H_1'$ with $\boldsymbol{\theta}' = (\zeta, \sigma^2)$ where $\zeta = \frac{\mu}{\sigma}$. The joint prior is given by eq. S14 as

$$p(\boldsymbol{\theta}') = \frac{\beta^\alpha}{\Gamma(\alpha)}(\sigma^2)^{-\alpha-1} \mathrm{e}^{-\beta/\sigma^2} \cdot \frac{1}{\sqrt{2\pi k}} \mathrm{e}^{-\frac{1}{2k}\zeta^2}, \quad \text{(S18)}$$

which gives the conditional prior

$$p(\sigma^2|\zeta = 0) \propto (\sigma^2)^{-\alpha-1} \mathrm{e}^{-\beta/\sigma^2}. \quad \text{(S19)}$$

This is $\mathrm{IG}(\alpha, \beta)$ and matches the prior under $H_0$.

Thus conditioning on $\mu = 0$ and on $\zeta = \frac{\mu}{\sigma} = 0$ induces different distributions of $\sigma^2$. Figure S1A&B show the $\mu$-$\sigma^2$ plane for the prior $p(\mu, \sigma^2)$, and large values of $\sigma^2$ are clearly more common in the shaded area in B than in the shaded area in A. The conditioning event, $\zeta = \frac{\mu}{\sigma} = 0$, is somewhat informative about $\sigma^2$, implying that $\sigma^2$ tends to be large.

To apply the S-D density ratio using $\zeta$, note that the prior is $\pi(\zeta = 0) = (2\pi k)^{-1/2}$ and the posterior is

$$\pi(\zeta = 0|\boldsymbol{x}) = \sqrt{\frac{1+nk}{2\pi k}} \left[\frac{2\beta + ns^2 + n\bar{x}^2}{2\beta + ns^2 + \frac{n\bar{x}^2}{1+nk}}\right]^{-\alpha-\frac{n}{2}} \quad \text{(S20)}$$

under $H_1$ (see Examples 1.6 & 7.3 in O'Hagan and Forster, 2004), so that the S-D ratio $B_\zeta = \frac{\pi(\zeta=0)}{\pi(\zeta=0|\boldsymbol{x})}$ gives $B$ of eq. S16, as expected.

In contrast, with parameter $\mu$ in $H_1$, we have

$$\pi(\mu = 0) = (2\pi k\beta)^{-\frac{1}{2}} \cdot \frac{\Gamma(\alpha + \frac{1}{2})}{\Gamma(\alpha)}, \quad \text{(S21)}$$

and

$$\pi(\mu = 0|\boldsymbol{x}) = \left[1 + \frac{n^2k\bar{x}^2}{(1+nk)(2\beta + ns^2 + \frac{n\bar{x}^2}{1+nk})}\right]^{-\alpha-\frac{n}{2}} \times \left[\frac{k}{1+nk}(2\beta + ns^2 + n\bar{x}^2)\right]^{-\frac{1}{2}} \times \left[B\left(\frac{1}{2}, \alpha + \frac{n}{2}\right)\right]^{-1}. \quad \text{(S22)}$$

Thus the density ratio is

$$\begin{aligned} B_\mu &= \frac{\pi(\mu = 0)}{\pi(\mu = 0|\boldsymbol{x})} \\ &= \frac{1}{\sqrt{1+nk}} \left[1 + \frac{n^2k\bar{x}^2}{(1+nk)(2\beta + ns^2 + \frac{n\bar{x}^2}{1+nk})}\right]^{\alpha+\frac{n}{2}} \\ &\quad \times \sqrt{\frac{2\beta + ns^2 + n\bar{x}^2}{2\beta}} \cdot \frac{\Gamma(\alpha + \frac{1}{2})}{\Gamma(\alpha)} \frac{\Gamma(\alpha + \frac{n}{2})}{\Gamma(\alpha + \frac{n+1}{2})} \\ &= B \times c, \end{aligned} \quad \text{(S23)}$$

where $c$ is the second factor in the product. Thus due to the mismatch of priors between models, $p(\sigma^2|H_0) \neq p(\sigma^2|\mu = 0, H_1)$, $B_\mu$ differs from the Bayes factor $B$ (eq. S16) by the factor $c$.

### References


O'Hagan, A. and Forster, J. 2004. *Kendall's Advanced Theory of Statistics: Bayesian Inference*. Arnold, London.

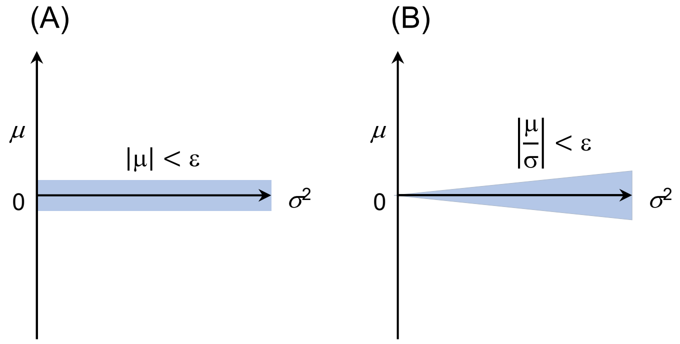


**Fig. S1**: Illustration of the Borel-Kolmogorov paradox. Given the joint distribution $p(\mu, \sigma^2)$, $-\infty < \mu < \infty, \sigma^2 < \infty$, conditioning (A) on $\mu = 0$ and (B) on $\frac{\mu}{\sigma} = 0$ may lead to different conditional distributions for $\sigma^2$, with $p(\sigma^2|\mu = 0) \neq p\left(\sigma^2\middle|\frac{\mu}{\sigma} = 0\right)$. Note that in each case the conditional distribution is the limit of the distribution of $\sigma^2$ over the shaded area, when the size of the shaded area approaches zero (i.e., when $\epsilon \longrightarrow 0$).

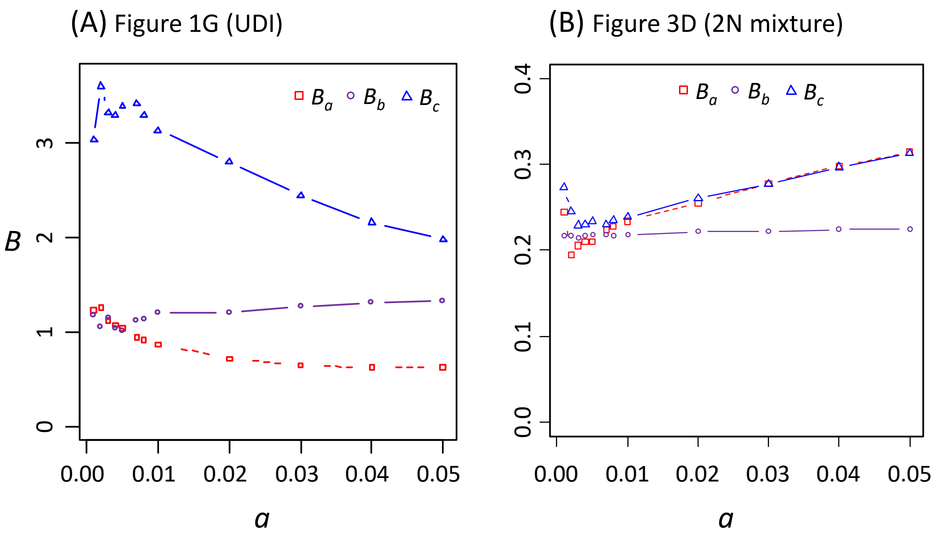


**Fig. S2**: (A) Bayes factor for testing gene flow under the UDI model calculated using the S-D density ratio applied to the three segments $(a, b, c)$ for the data of Figure 1F&G, plotted against the factor $a$ for the prior standard deviation, so that $\epsilon_\varphi = \epsilon_\beta = a\frac{1}{\sqrt{12}} = a \cdot 0.2886$, or $B_a = \frac{\epsilon_\varphi}{\mathbb{P}(\varphi<\epsilon_\varphi|\boldsymbol{x})}$, $B_b = \frac{\epsilon_\beta}{\mathbb{P}(1-\beta<\epsilon_\beta|\boldsymbol{x})}$, and $B_c = \frac{\epsilon_\varphi}{\mathbb{P}(1-\varphi<\epsilon_\varphi|\boldsymbol{x})}$, where $\beta = \tau_Y/\tau_R$. (B) Bayes factor for testing Gaussian mixture (eq. 29) for the data of figure 3D, plotted against the factor $a$, with the differentials given as $\epsilon_\alpha = a/\sqrt{12}$ and $\epsilon_\mu = a$.

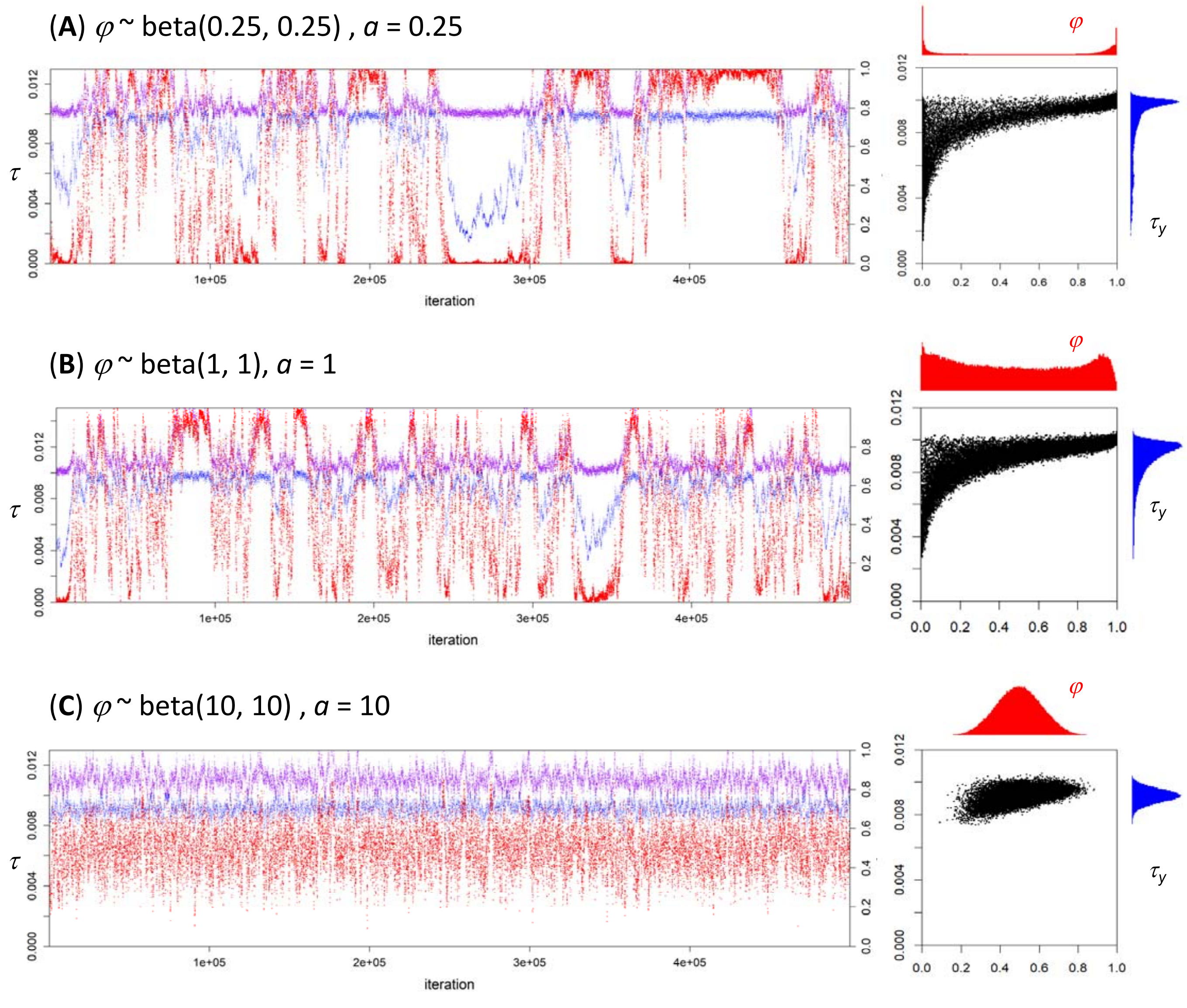


**Fig. S3**: Trace plots and scattergrams for parameters under the UDI model (purple for $\tau_R$, blue for $\tau_Y$ and red for $\varphi$ (the 2nd $y$-axis) in analysis of the data of figure 1F&G under three priors, $\varphi \sim \text{Beta}(a, a)$, with (**A**) $a = \frac{1}{4}$, (**B**) $a = 1$, and (**C**) $\alpha = 10$. Panel (**B**) is from figure 1. See legend to figure 1.

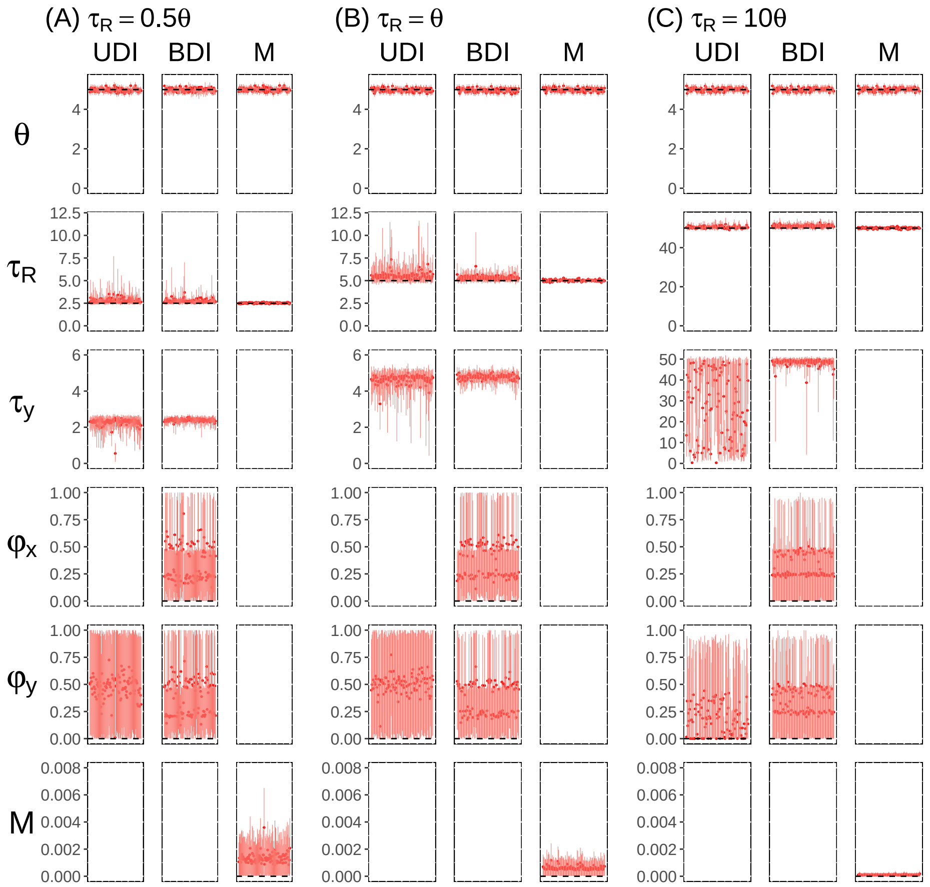


**Fig. S4**: Posterior means and 95% HPD CIs of parameters in BPP analysis of 100 replicate datasets simulated under $H_0$: MSC (no gene flow) and analyzed under the MSC-I and MSC-M models. For MSC-I, both the UDI and BDI models are used. Results of Bayesian test of gene flow using those data are in table 3.

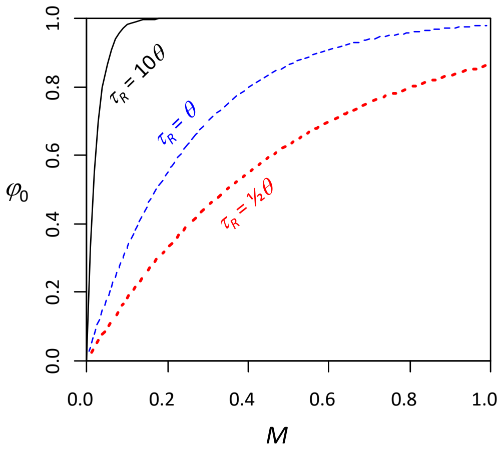


**Fig. S5**: The expected proportion of introgression under the MSC-M model ($\varphi_0$, eq. 37).
